# The differential effect of SARS-COV-2 NSP1 on mRNA translation and stability reveals new insights linking ribosome recruitment, codon usage and virus evolution

**DOI:** 10.1101/2024.06.17.599344

**Authors:** Juan José Berlanga, Tania Matamoros, Miguel Rodríguez Pulido, Margarita Sáiz, Mercedes Núñez Bayón, René Toribio, Iván Ventoso

## Abstract

The non-structural protein 1 (NSP1) of SARS-CoV-2 blocks the mRNA entry channel of the 40S ribosomal subunit, causing inhibition of translation initiation and subsequent degradation of host mRNAs. However, target mRNA specificity and the way in which viral mRNAs escape NSP1-mediated degradation have not been clarified to date. Here we found that NSP1 acts as a translational switch capable of blocking or enhancing translation depending on how preinitiation complex 43S-PIC is recruited to the mRNA, whereas NSP1-mediated mRNA degradation mostly depends on codon usage bias. Thus, fast-translating mRNAs with optimal codon usage for human cells that preferentially recruit 43S-PIC by threading, showed a dramatic sensitivity to NSP1. On the contrary, slow-translating mRNAs with suboptimal codon usage and 5’ UTR that enabled slotting on 43S-PIC were resistant to or even enhanced by NSP1. Translation of SARS-CoV-2 mRNAs escapes NSP1-mediated inhibition by a proper combination of suboptimal codon usage and slotting-prone 5’ UTR that also confers efficient translation. Thus, the prevalence of non-optimal codons found in SARS-CoV-2 and other coronavirus genomes is favored by the distinctive effect that NSP1 plays on translation and mRNA stability.

## Introduction

Translation in eukaryotes not only regulates protein outputs but also mRNA stability. Cap-dependent translation initiates with the recruitment of the 40S ribosomal subunit to the 5’ cap of mRNA, and the subsequent scanning of 43S-PIC through the 5’ UTR to locate the initiation codon ^1–3^. Binding of the eIF4F complex (eIF4E+eIF4G+eIF4A) to the 5’ cap promotes local unwinding of the mRNA secondary structure that facilitates attachment of 43S-PIC ^4^. Features of the 5’ UTR including length, nucleotide composition and the presence of stable secondary structures can greatly influence translation initiation rates of particular mRNAs ^3–6^. Thus, initiation on mRNAs with a complex 5’ UTR is more dependent on the helicase activities of eIF4A and DDX3, which are associated to scanning 43S-PIC, whereas translation of mRNAs with an unstructured 5’ UTR usually shows fewer requirements for initiation ^7–11^. Although structural models of 43S-PIC bound to short mRNAs are available (48S-PIC), the way mRNA is inserted into the 40S subunit for codon inspection is still a matter of debate ^1^. Threading into the mRNA entry channel or lateral slotting on the groove of the 40S subunit have both been proposed ^12–15^. Threading and slotting would show different requirements for 5’ UTR length and topology, raising the possibility that both mechanisms may operate depending on the 5’ UTR features found in a particular mRNA.

Translation elongation is increasingly recognized for its role in regulating mRNA stability and protein quality control, but it can also influence translation initiation by a yet unknown mechanism ^16–19^. mRNAs with a synonymous codon composition adapted to aa-tRNA availability in a given cell type or organism tend to be more stable, and they are translated faster ^19^. Since optimal codons usually end in C or G, mRNAs with optimal codon usage also show a higher G+C content. Although this may result in a more stable secondary structure that could hinder ribosome advance, mRNAs with optimal codons show an effective ribosome flow that minimizes ribosome stalling-mediated mRNA destabilization^17,19^.

In stark contrast, many viral genomes including SARS-CoV-2 show a low G+C content and a suboptimal codon usage for human cells, mainly shaped by a persistent C-to-U mutational bias ^20–23^. Despite this, SARS-CoV-2 mRNAs are efficiently translated in a context of generalized translation shut-off and host mRNA degradation induced by the viral non-structural protein 1 (NSP1) ^24–28^. NSP1 of SARS-CoV-2 binds with high affinity to the 40S ribosomal subunit at the mRNA entry channel, thus preventing penetration of mRNA through this region ^29–31^. This translational blockade by NSP1 results in a generalized host mRNA degradation ^26–28,32^. While the C-terminal domain of NSP1 is inserted into the mRNA entry channel, the remaining N-terminal domain stabilizes the NSP1-40S complex by interacting with surrounding ribosomal proteins (e.g. RPS3) and some initiation factors (e.g. eIF3G) ^29,33–36^. More recently, a dynamic binding of N-terminal domain of NSP1 to the decoding center of 40S has been described ^24^. Although most attempts failed to detect NSP1-associated nuclease activity, the N-terminal domain was shown to be essential for NSP1-mediated mRNA degradation, also requiring the presence of eIF3G and eIF4F in the PIC ^34,36^. Another unresolved question is how SARS-Cov-2 mRNAs escape the inhibitory activity of NSP1. The presence of a cap-proximal stem-loop (SL1) in the 5’ leader (5’ L) of genomic and subgenomic SARS-CoV-2 mRNAs has been reported as essential to escape the inhibitory effect of NSP1 ^25,34,35,37–39^. Although a proven mechanistic model for this resistance is still lacking, some authors have proposed that binding of SL1 to N-terminal domain would displace NSP1 from the mRNA entry channel, allowing the attachment of viral 5’ L to 43S-PIC ^40^. However, a direct interaction of NSP1 with 5’ L was not observed in other reports ^34^.

Here, we have analyzed the contribution of the 5’ UTR and codon usage bias to the sensitivity of mRNAs to NSP1 in human cells. Our results reveal new insights and links between 43S-PIC recruitment to mRNA, codon usage bias and viral evolution.

## Results

### Differential effect of NSP1 on reporter mRNA translation and stability

HEK293T cells were co-transfected with increasing amounts of NSP1-expressing plasmid in combination with several reporter genes available in our lab (Figure 1A). Each reporter was expressed from the same plasmid, and the 5’ UTRs of the resulting mRNAs were nearly identical, consisting of 82-94 nt with a 60% G+C and moderate secondary structure (5’ UTR 82/94) (Figure S1A). The transcription initiation site for these constructs was confirmed by 5’ RACE (Figure S1B). We found that the levels of NSP1 WT accumulated in transfected cells were comparable to those found in SARS-CoV-2-infected cells (Figure S1C). Furthermore, NSP1 was only present in the 40S/PIC fractions, as described in some previous reports (Figure S1D) ^29,32^. Whereas the expression of some reporter genes such as dsRed, EGFP and NanoLuc was dramatically inhibited by NSP1 (up to 30-fold), the expression of Firefly (FFL) and Renilla (RL) luciferases was much less affected at any of the NSP1 doses tested (Figure 1A). To determine whether this inhibitory effect was specific to NSP1, we analyzed the impact of other translation initiation inhibitors such as silvestrol and oligo 4-VIC, which block eIF4A helicase activity and the ES6S region of 40S, respectively ^14,41,42^. Despite the proximity of the binding sites of all three inhibitors to the mRNA entry channel of the 40S subunit, oligo 4-VIC and silvestrol similarly inhibited the expression of all reporter genes tested (Figure 1B), suggesting that the differential effect of NSP1 was not the simple result of translation inhibition. Since NSP1 binding to the ribosome not only blocks translation but also induces mRNA degradation, we compared the effect of wild-type (WT) NSP1 with previously described mutants deficient in ribosome binding (K164A/H165A, M1) or lacking the mRNA degradation activity (R124A/K125A, M2) ^34,36^. Interestingly, RL and FFL expression showed a similar sensitivity to NSP1 WT and M2, whereas the M2 mutant had a much lower inhibition of EGFP, dsRed and NanoLuc compared with NSP1 WT (Figure 1C). To confirm whether RL and FFL mRNAs were resistant to NSP1-induced degradation, we quantified the accumulation of reporter mRNA in transfected cells. Indeed, NSP1 WT induced a strong reduction of EGFP, dsRed and NanoLuc mRNAs, but not of RL and FFL mRNAs (Figure 1D).

**Figure 1.**
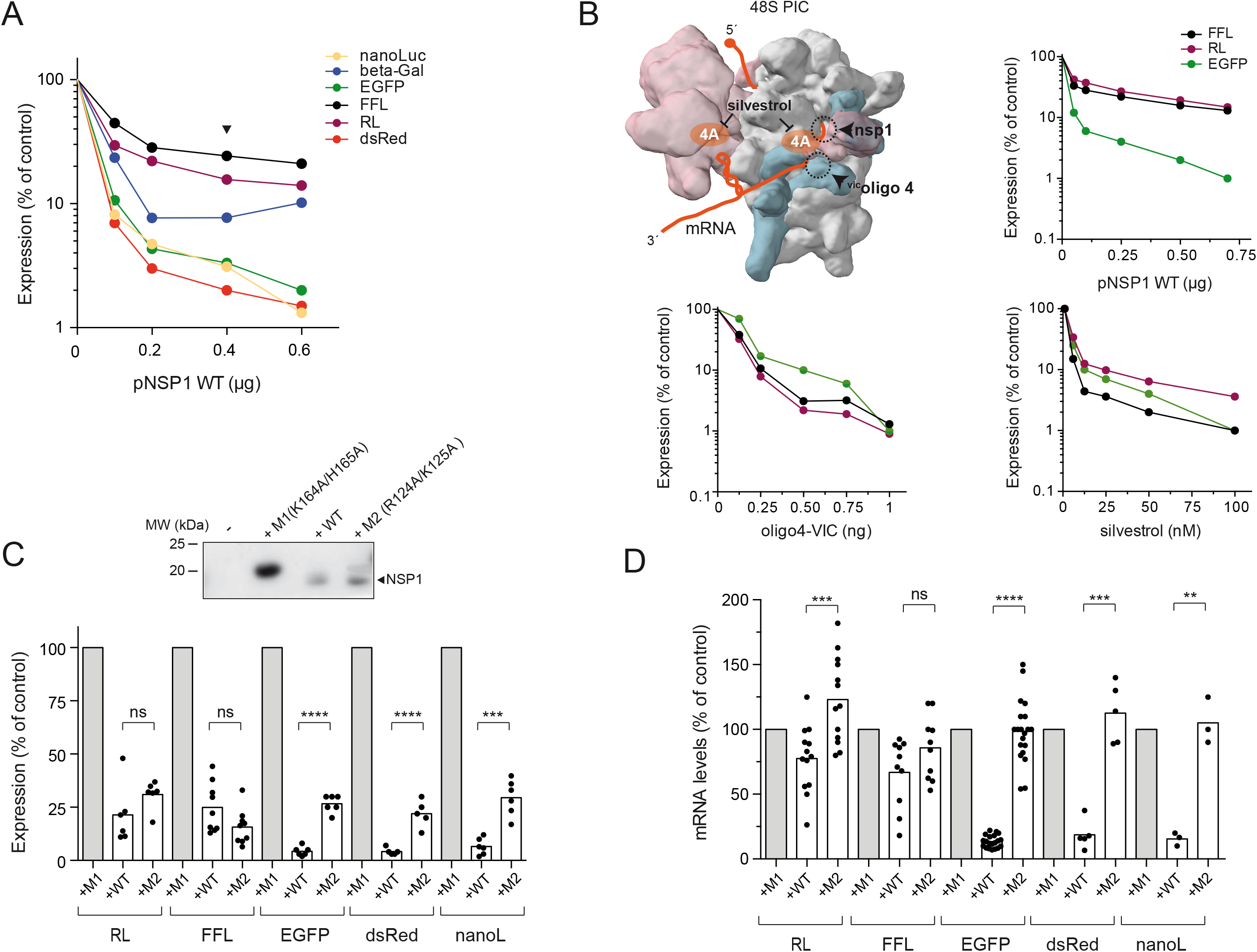
Differential sensitivity of reporter gene expression to NSP1. (A) Effect of increasing amount of NSP1 WT-expressing plasmid on expression of the indicated reporter genes in HEK293T cells. Data are the average of two independent experiments and are expressed as a percentage relative to the cells transfected with 0.6 μg of pNSP1 M1 (*null* mutant, control). Arrow indicates the amount of pNSP1 plasmid selected for further experiments. (B) Comparative analysis of the effect of NSP1 WT, oligo4^VIC^ and silvestrol on expression of FFL, RL and EGFP reporters. The binding sites of these inhibitors on 48S-PIC model are indicated (some eIFs were omitted for simplicity). Data are the average of two independent experiments. (C) Comparative analysis of the effect of NSP1 WT and M2 (R124A/K125A, mRNA degradation *null*) on the expression of the indicated reporter genes. A representative western blot monitoring the expression of different NSP1 variants is also shown. (D) Comparative analysis of the effect of NSP1 WT and M2 on accumulation of the indicated reporter mRNA 24 h after transfection. mRNA levels were quantified by RT-qPCR as described in Materials and Methods. Significance was scored by two-tailed, unpaired *t*-test as described in Materials and Methods.

### Codon usage of CDS mRNA influences sensitivity to NSP1

Since the reporter genes tested are transcribed from the same promoter (CMV) and contain almost identical 5’ and 3’ UTR regions flanking the coding sequences (CDS), we analyzed the correlation between sensitivity to NSP1 and a number of CDS parameters including codon composition (usage), G+C content and predicted RNA secondary structure (Figures 2A and S2A). To strengthen the analysis, we included additional reporter genes of cellular (e.g., human actin (ACTB)) and viral origin (e.g., nucleocapsid (N) from SARS-CoV-2) that showed different codon usage and nucleotide composition (Supplementary table 1). As a measure of codon bias, we used codon adaptation index (CAI), tRNA adaptation index (tAIg) and codon stability coefficients (CSCg) estimated for HEK293T cells ^43,44^. Although based on different parameters, all three metrics are related and can be used as predictors of codon bias and expression level ^45^. Interestingly, we found a positive and strong correlation between sensitivity to NSP1 WT and optimal codon usage of the reporter CDS for each of the three metrics used (Figure 2A). Importantly, no correlation was found for NSP1 M2, suggesting that codon composition of reporter genes may also influence NSP1 WT-induced mRNA degradation (Figure 2A). A similar positive correlation was found between sensitivity to NSP1 WT and G+C content, both analyzed globally and specifically at the third-codon position (GC3) (Figure S2A). This result was expected since both G+C and GC3 contents are linked to codon bias ^46^.

**Figure 2.**
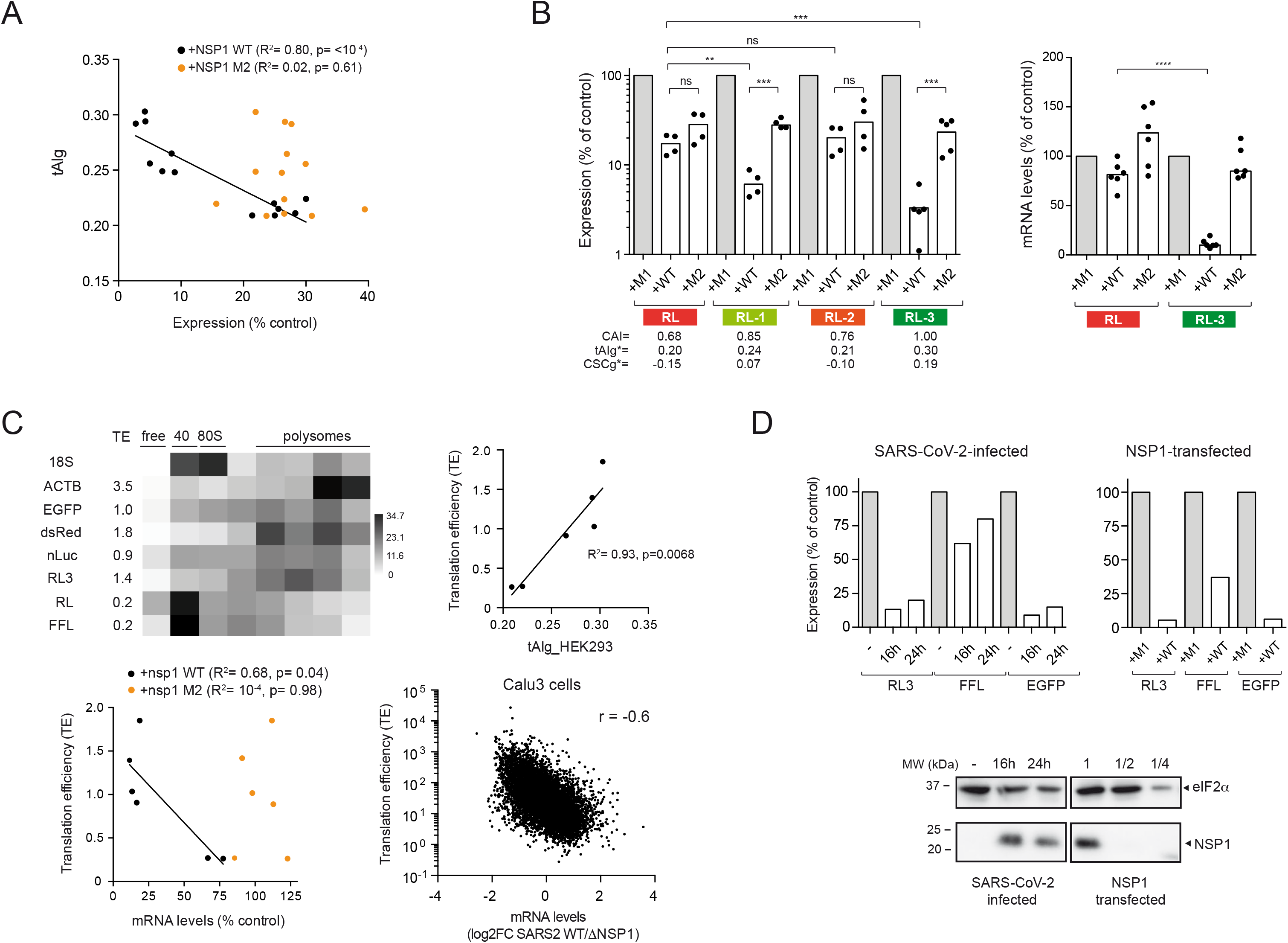
Influence of mRNA codon usage bias on sensitivity to NSP1. (A) Correlation between tRNA adaptation index (tAIg) of reporter mRNAs and their corresponding sensitivities to NSP1 WT or M2. Data were scored as described in Figure 1. The R^2^ and p-values are indicated. (B) Effect of RL CDS recoding on sensitivity to NSP1 WT and M2. The CAI, tAIg and CSCg values for parental (RL red) and RL-1 to 3 variants (orange to green) are indicated. Right panel shows a relative quantification of RL and RL-3 mRNA levels in cells expressing NSP1 WT or M2. (C) Correlation between translation efficiency (TE), tAIg and sensitivity to NSP1 WT and M2. Upper left panel shows TE calculated for the indicated mRNAs by polysome profiling followed of RT-qPCR. The resulting heat map of the relative quantification is shown. Correlation between TE and tAIg (upper right panel) or TE and sensitivity to mRNA-induced degradation by NSP1 WT or M2 (bottom panel). Data were scored as described above and the resulting R^2^ is indicated. Bottom right panel shows the correlation between TE of human mRNAs and sensitivity to NSP1-induced mRNA degradation in Calu3 cells infected with SARS-CoV-2 WT compared with that infected by a mutant virus defective in NSP1 (⊗NSP1). Data were taken from GSE200422 ^27^ and analyzed as described in Materials and Methods; the resulting Spearman correlation coefficient (r) is indicated. (D) Effect of SARS-CoV-2 infection on expression of RL-3, FFL and EGFP in VeroE6 cells. Cells were infected, transfected with a combination of the three plasmids and analyzed at 16 and 24 hpi for luciferase activity (upper panel) or by western blot against NSP1, RPS6 and EGFP. In parallel wells, cells were co-transfected with pNSP1 and a combination of RL-3, FFL and EGFP plasmids and analyzed 20 h later (top panels). A comparative analysis of NSP1 levels accumulated in infected and transfected cells is shown (bottom panel). Blots were also probed with anti-eIF2α as loading control. The 1/2 and 1/4 dilutions of cell extracts are also indicated.

To test the influence of codon bias on sensitivity to NSP1 WT, we recoded the RL and EGFP CDS to generate variants with different codon usage. From the parental RL, which shows a suboptimal codon usage (tAIg= 0.2), we generated two variants with increased codon adaptation index to HEK293T cells (Figure 2B). The resulting codon-optimized RL-1 and RL-3 constructs showed an increased expression compared to the parental RL (Figure S2B), and a dramatic increase in sensitivity to NSP1 WT (up to 30-fold for RL-3 variant), but not to NSP1 M2, suggesting that the corresponding codon-optimized RL variants became more sensitive to NSP1-mediated mRNA degradation (Figure 2B and S2B). To confirm this, we compared the amount of RL and RL-3 mRNA accumulating in cells expressing NSP1s, revealing a strong reduction of RL-3 mRNA in the presence WT NSP1 (Figure 2B). Conversely, we de-optimized the parental EGFP (tAIg= 0.3) to generate two variants (EGFP-1 and 2) with a predicted suboptimal codon usage for HEK293T cells (Figure S2C). Notably, codon de-optimization of EGFP drastically reduced the sensitivity to NSP1 WT from 30-fold to 2-3-fold inhibition, a rate that was similar to the parental RL and FFL reporters (Figure S2C). These results indicate that genes with optimal codon usage are more sensitive to NSP1-induced inhibition.

Since optimal codon usage accelerates the rate of translation elongation, we measured the translation efficiency (TE) of some reporter mRNAs by polysome profiling. As expected, reporter mRNAs with optimal tAIg (dsRed, EGFP, NanoLuc, RL3) showed a higher TE, and conversely, reporter mRNAs with lower tAIg values (FFL and RL) accumulated preferentially in monosome and 40S fractions, which is indicative of a slower rate of translation initiation (Figure 2C). We found a strong direct correlation between TE and sensitivity to NSP1 when analyzed at both protein (reporter activity) and mRNA levels (Figure 2C). To further confirm this, we used previously published data to calculate TE of human mRNAs in Calu3 cells and the impact of SARS-CoV-2 WT infection on mRNA levels ^27^. Similar to that found with reporter mRNAs, a significant correlation was observed between TE and NSP1-mediated degradation of human mRNAs; the mRNAs that were more sensitive to NSP1 in SARS-CoV-2-infected cells also showed higher translation rates in uninfected cells (Figure 2C).

To experimentally confirm these findings in SARS-CoV-2-infected cells, we compared the expression of EGFP, RL-3 and FFL in VeroE6 cells, infected or not with SARS-CoV-2. Notably, expression of RL3 and EGFP was more sensitive than FFL to SARS-CoV-2 infection, although the observed inhibition was less pronounced than that found in VeroE6 cells transfected with pNSP1 plasmid (which accumulated ∼2-fold more NSP1 than in infected cells) (Figure 2D). Therefore, the differential effect of NSP1 on reporter genes with different codon usages and TE was also seen in SARS-CoV-2-infected cells.

### Widely diverse influence of 5’ UTR on NSP1-mediated effect

Next, we systematically analyzed the influence of 5’ UTR on sensitivity to NSP1 using a collection of parental RL and FFL constructs bearing different 5’ UTRs (Figure 3A and supplementary table 2). We analyzed the influence of 5’ UTR length (from 8 to 650 nt), the presence of stable secondary structures at different positions of the 5’ UTR and the influence of nucleotide composition and motifs in the 5’ UTR. We found great diversity in effects of 5’ UTRs on NSP1-mediated effect, ranging from those that conferred an extreme sensitivity (greater than 40-fold inhibition) to those that allowed enhanced expression of the reporter gene in the presence of NSP1 WT. Regarding the influence of 5’ UTR length, a progressive increase in length up to 650 nt strongly reduced the expression of both RL and FFL constructs, but had a minor effect on sensitivity to NSP1 (Figure 3A). However, shortening the 5’ UTR up to 5 nt in an RL construct bearing the TISU element (TISU-RL) dramatically increased sensitivity to NSP1 WT, and to a lesser extent to NSP1 M2 (Figure 3A and B). TISU provides an optimal AUG context to promote efficient translation of mRNA bearing extremely short 5’ UTRs ^47,48^. We confirmed this effect by placing the TISU element upstream of EGFP CDS, resulting in an almost complete inhibition of EGFP expression by NSP1 WT (Figure 3B). The levels of TISU-containing mRNAs were also further reduced by NSP1 WT compared to parental constructs, especially for TISU-EGFP mRNA that dropped 15-fold in the presence of NSP1 (Figure 3B).

**Figure 3.**
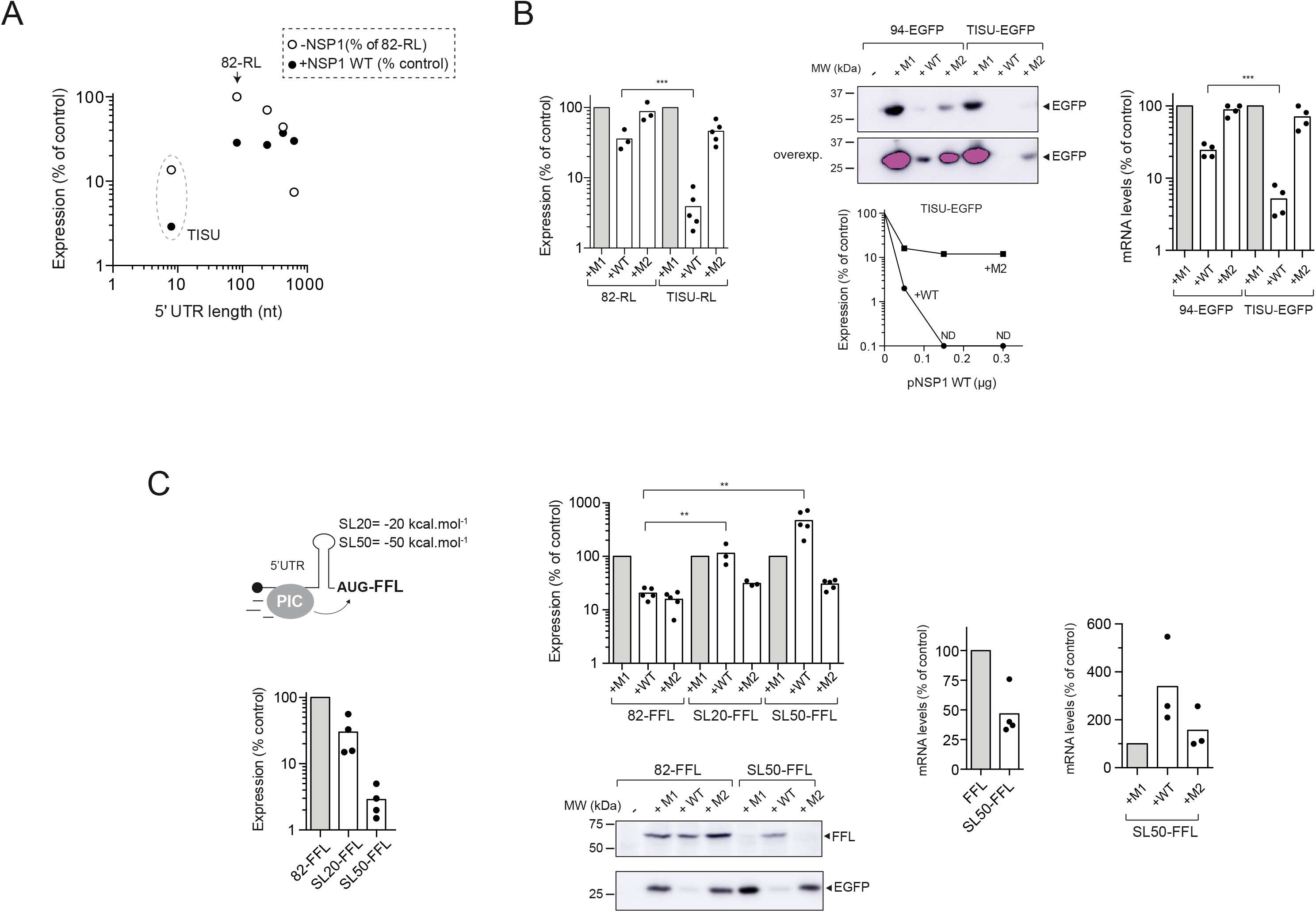
Influence of 5’ UTR length and secondary structure on sensitivity to NSP1. (A) RL plasmids with 5’ UTRs ranging from 8 to 650 nt were co-transfected with FFL plasmid and the resulting normalized activities were expressed as a percentage relative to the 82-RL construct (-NSP1). The sensitivity to NSP1 WT expression (+NSP1) for every construct is expressed as described in previous figures. (B) Effect of NSP1 WT and M2 on 82-RL and TISU-RL expression (left panel) or 94-EGFP and TISU-EGFP expression assayed by western blot (middle panel). The effect of increasing amount of pNSP1 WT or M2 plasmids on TISU-EGFP expression is also shown. ND, not detected. Right panel shows mRNA level quantification of 94-EGFP and TISU-EGFP in cells expressing NSP1 WT or M2. Significance was scored as described before. (C) Influence of stable stem-loops (SLs) placed immediately before the AUG of FFL on the sensitivity to NSP1s. The placement and stability (⊗G°) of SLs are shown (left panel). Effect of SLs on FFL expression relative to 82-FFL (lower left panel) and sensitivity to NSP1 WT and M2 (top middle panel). SL50-FFL expression relief by NSP1 WT was also confirmed by western blot using anti-Luc antibodies (bottom middle panel). Right panels shows relative quantification of 82-FFL and SL50-FFL mRNA levels in control and cells co-expressing NSP1 WT or M2.

Next, we analyzed the contribution of stable secondary structures in the 5’ UTR to NSP1-mediated inhibition. It is well known that stable stem loops (ΔG° <-20 kcal.mol^-1^) can delay or even prevent scanning of PIC-43S on mRNA during translation initiation ^5,6^. We used previously described constructs generated from parental 82-FFL bearing a stem loop of -20 (SL20-FFL) or -50 kCal.mol^-1^ (SL50-FFL) immediately upstream of the AUGi. The resulting constructs showed a reduced expression, especially for SL50 that was 30- to 50-fold lower than parental 82-FFL (Figure 3C). Moreover, we also observed reduced levels of SL50-FFL mRNA compared with 82-FFL, suggesting that SL50-induced translation blockade resulted in some mRNA destabilization (Figure 3C). Surprisingly, the expression of SL50-FFL was stimulated ≈ old by NSP1 WT, but not by NSP1 M2 (Figure 3C). We also found that NSP1 WT increased the levels of SL50-FFL mRNAs, suggesting that NSP1-mediated relief of SL50-FFL mRNA translation can result in mRNA stabilization (Figure 3C). A similar result was observed when SL50 was placed in RL, although in this case the stimulation by NSP1 WT co-expression was only ≈2-fold (Figure S3A and B). We also confirmed by western blot that the size of the luciferase band accumulated in NSP1-expressing cells was similar to that of control samples, showing that NSP1-mediated bypass of SL50 involves initiation at the natural AUGi of FFL mRNA (Figure 3C).

### G-rich eIF4A binding motifs proximal to 5’ cap determines sensitivity to NSP1

We next analyzed the influence of nucleotide composition of the 5’ UTR on sensitivity to NSP1 (Figure 4A). We used a previously described RL construct bearing a 5’ UTR lacking Gs (G-less-RL) except the one located in the second position from the transcription start site (TSS), which is necessary for transcription from the CMV promoter ^14^. G-less 5’ UTR reduced the expression of parental RL, but it had little effect on the expression of codon-optimized RL3 (Figure 4B). Interestingly, the presence of NSP1 WT but not M2, enhanced the expression of G-less-RL by up to 10-fold (Figure 4B). We also observed a slight increase in mRNA levels of G-less-RL in the presence of NSP1 WT, suggesting that translational relief by NSP1 can result in some mRNA stabilization (Figure S4A). In addition, the presence of G-less 5’ UTR made RL3 resistant to NSP1 WT (Figure 4B). The effect of NSP1 on the expression of some of these constructs was also confirmed in other cell types including HeLa, MEF and Calu-3 cells (Figure S4B). Similar results were obtained when 5’ UTR-82 of EGFP and FFL was replaced by a 5’ UTR low in Gs (G-low), unrelated in sequence to G-less (Figures S4C and D). These results showed that NSP1 WT can act as a translation enhancer of mRNAs bearing G-poor 5’ UTR and suboptimal codon usage.

**Figure 4.**
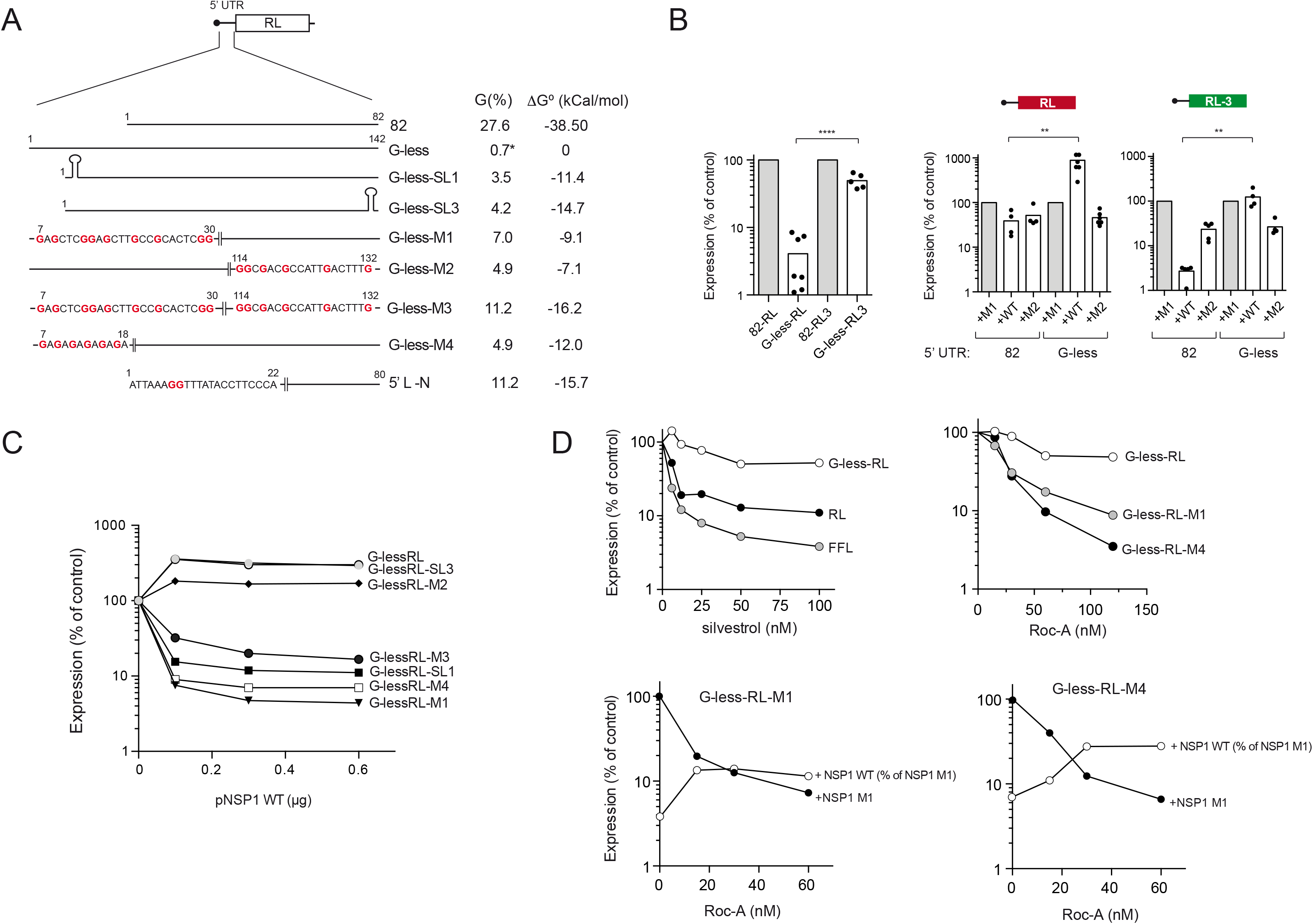
The presence of Gs near the 5’ cap critically determines the sensitivity to NSP1. (A) Schematic diagram of G-less 5’ UTR and its variants including G, SL or GA motifs at the indicated positions. The composition (% G) and stability (ΔG° in kcal/mol) of the resulting sequences are indicated. The Gs introduced in the sequences are in bold. The first 22-nt sequence of 5’ L N of SARS-CoV-2 is also shown. (B) Left panel shows a comparative analysis of RL expression with G-less or 82 5’ UTRs in combination with optimal or suboptimal codon usage (RL vs RL-3). Effect of NSP1 WT or M2 on the expression of G-less 5’ UTR in combination with optimal or suboptimal codon usage (middle and right panels). Significance was scored as described in previous figures. (C) Sensitivity of G-less and its variants to increasing amounts of NSP1 WT. Data are the average of two independent experiments. (D) Reduced sensitivity of G-less-RL expression to eIF4A inhibition by silvestrol (top left panel) or RocA (top right panel) compared to RL, FFL or M1 and M4 variants of G-less-RL. Inhibitors were added 1 h after transfection and measurements were taken 16 h later. Bottom panels show the residual expression of G-less-RL-M1 and M4 in the presence of NSP1 WT for every RocA concentration. Data are expressed as % respect to samples with NSP1 M1 and the corresponding RocA concentration.

We next introduced Gs and/or secondary structure in G-less 5’ UTR distally or proximally to 5’ cap (Figure 4A). Whereas introduction of Gs or a stem loop of moderate structure immediately before AUGi did not affect NSP1 WT-mediated stimulation, introduction of 9 Gs spaced by 1-5 nt near the 5’ cap (G-less-M1) drastically increased sensitivity to NSP1. Thus, expression of G-less-M1 was inhibited by more than 10-fold, resulting in a net change of almost 100- fold when compared with G-less (Figure 4C). Introduction of a stem loop of moderate stability also made G-less sensitive to NSP1 WT-mediated inhibition, although the change was less dramatic (Figure 4C). This result suggests that the presence of Gs near the 5’ cap had a greater impact on sensitivity to NSP1 than a tendency to form secondary structures. Since helicase eIF4A binds purine rich motifs in the 5’ UTR of many mRNAs ^49,50^, we inserted a (GA)_6_ motif near the 5’ cap (G-less-M4) to test the role of eIF4A in the susceptibility to NSP1. Similar to G-less-M1, expression of G-less-M4 construct was strongly inhibited by NSP1, suggesting that eIF4A binding proximal to mRNA 5’ cap could determine the sensitivity to NSP1. To confirm this, we compared the sensitivity of G-less-RL constructs to silvestrol and RocA, two inhibitors that clamp eIF4A on target mRNAs ^49^. Notably, we found that G-less-RL expression was much less sensitive to eIF4A inhibitors than reporter genes carrying the 5’ UTR-82, thus confirming our previous *in vitro* data ^14^. Similar results were found when the sensitivity of G-low-FFL to RocA was compared to parental 82-FFL (Figure S4E). On the contrary, expression of G-less-M4 and G-less-M1 showed an increased sensitivity to RocA similar or even superior to 82-RL (Figure 4D). Interestingly, we also found that the residual expression of G-less-M1-RL and G-less-M4-RL in the presence of NSP1 WT showed a lower sensitivity to RocA, suggesting that NSP1 allowed some eIF4A-independent translation initiation under these circumstances (Figure 4D).

### NSP1 blocks threading but allows slotting of mRNA into 43S-PIC

The differential effect observed on translation suggested that NSP1 could be blocking the threading of some mRNAs (e.g. TISU and G-less-M1/M4), while allowing the slotting of 43S-PIC on other mRNAs (e.g. G-less/G-low). To further substantiate this idea, we first progressively shortened parental 5’ UTR G-less from 142 nt to 8 nt, thereby reducing the chance of 43S-PIC slotting on the resulting 5’ UTR. Whereas NSP1 was still able to enhance the expression of 22 nt-long G-less 5’ UTR, and to a lesser extent of 15 nt-long G-less, a further reduction to 12 nt abruptly changed the effect of NSP1 on expression, and reached a 30-fold inhibition when the 5’ UTR was shorted to 8 nt (Figure 5A). Since a full insertion of mRNA into the 40S channel of 43S-PIC requires a minimum of 24 nt, the 12 nt-long G-less 5’ UTR was probably too short to allow a productive slotting in the presence of NSP1 even if some back scanning is considered ^12,15,42,51^. These results agree well with a blind spot of 10-30 nt from the 5’ cap detected when 48S-PIC was assembled *in vitro* on a 50 nt-long 5’ UTR mRNA, a result that has been taken as evidence of slotting ^12^. Since this 5’ UTR is very similar in composition to our G-less and G-low 5’ UTRs (see supplementary table 2), we inserted it in our RL plasmid (Slot-RL) and the resulting sensitivity to NSP1 was tested. Slot-RL showed a similar expression to G-less-RL that was resistant to NSP1 WT (Figure 5B).

**Figure 5.**
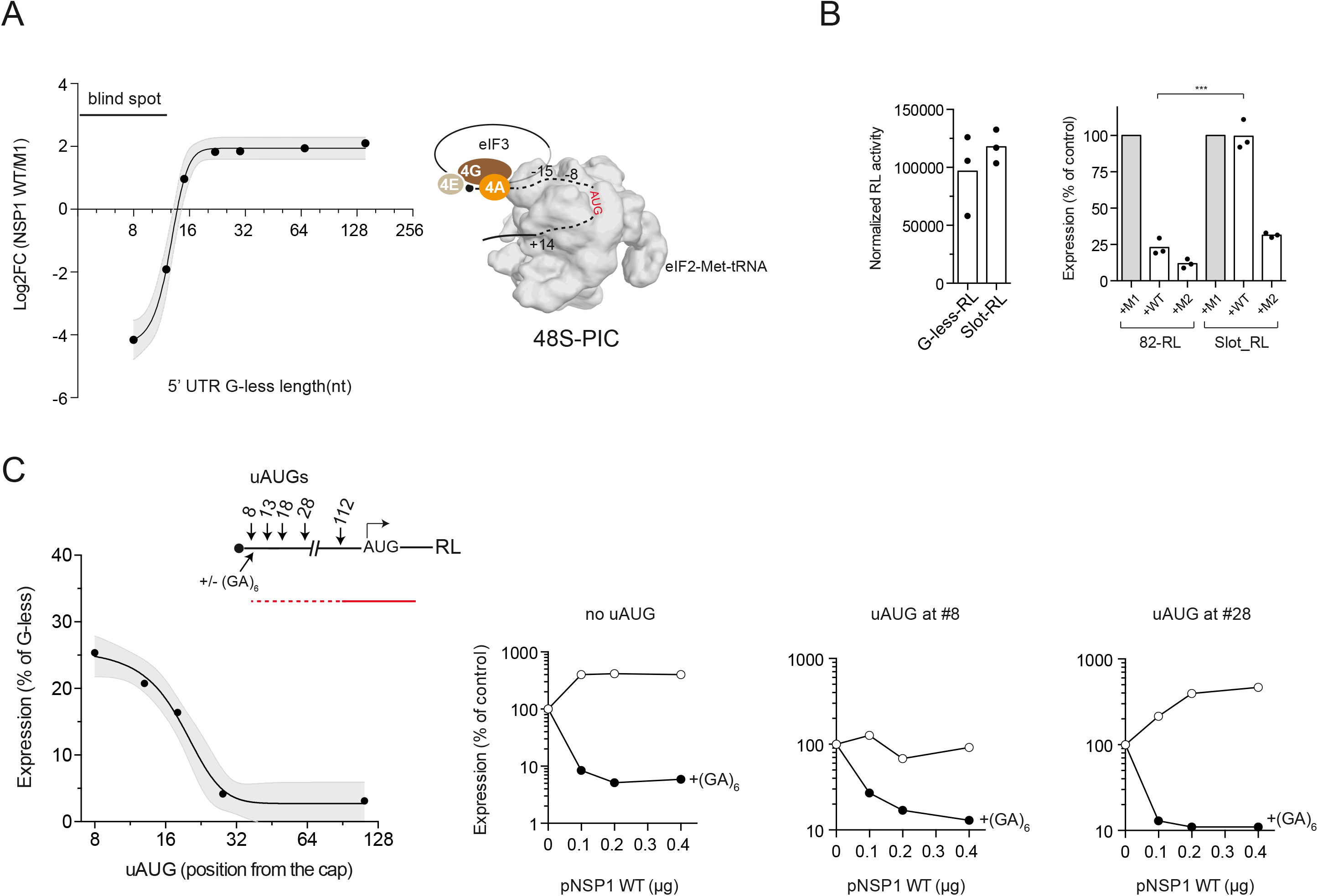
NSP1 promotes slotting of PIC to mRNA. (A) Effect of progressive shortening of 5’ UTR G-less on sensitivity to NSP1 WT. Co-transfections also included FFL plasmid to normalize for FFL activity. Data are expressed as log2 fold-change expression. Data points correspond to the mean from at least 3 independent experiments that were adjusted to a sigmoid function (R^2^= 0.92); gray shading represents the 95% confidence interval. The resulting estimated blind spot of 12-15 nt is shown. A model of 48S-PIC showing the position of mRNA residues along the mRNA channel of 40S is also represented (right panel). (B) Comparative analysis of the expression of G-less-RL and Slot-RL constructs (left panel). Comparative analysis of the sensitivity of 82-RL and Slot-RL to NSP1s (right panel). Significance was scored as described in previous figures. (C) Effect of upstream out-of-frame AUGs (uAUG) on expression of G-less-RL and sensitivity to NSP1. The uAUG was inserted where indicated and the resulting uORFs partially overlap the 5’ of the main RL ORF (red line). The presence of eIF4A-binding motif (GA)_6_ is indicated. HEK293T cells were cotransfected with the indicated constructs together with pFFL and the resulting expression was normalized for FFL activity and expressed as % of G-less-RL. Data represent the mean of four independent experiments that were adjusted to a sigmoid function (R^2^= 0.89); gray shading represents the 95% confidence interval (left panel). Middle and right panels show the effect of increasing amounts of pNSP1 WT plasmid on the expression of the indicated construct derived from pG-less-RL (open circles) or pG-less-M4 (black circles).

Next, we inserted upstream AUGs out of frame at 8, 13, 18, 28 and 112 nt from the 5’ cap in G-less-RL (Figure 5C). The resulting short uORFs overlap the first 25 nt of RL CDS to avoid the possibility of reinitiation, as occurs in ATF4 mRNA ^52^. Notably, whereas insertion of uAUG at positions 8 and 13 had only a moderate effect on expression (3-4-fold inhibition), the placement of uAUG at position 28 and onwards strongly reduced the expression of G-less-RL (25-30 fold) (Figure 5C), showing that a significant proportion of 43S-PIC can efficiently bypass any uAUG near the cap of G-less mRNA. Interestingly, the presence of NSP1 still enhanced the expression of G-less-uAUG_28, whereas the expression of G-less-uATG_8 was unaffected by NSP1 (Figure 5C). However, when (GA)_6_ motif was placed in these constructs (derived from G-less-M4-RL, see Figure 4A), a strong inhibition by NSP1 was observed (Figure 5C). These results showed that NSP1 allowed the slotting of 43S on G-less-RL mRNAs even beyond than expected according to the theoretical blind spot.

### Both 5’ UTR and codon usage of SARS-CoV-2 mRNAs promote resistance to NSP1

The next experiments were aimed at understanding how SARS-CoV-2 mRNAs can escape the inhibitory effect of NSP1. We first compared nucleotide composition and codon usage of full CDS of different coronavirus with those of human mRNAs (Figure 6A). As noticed before, a suppression in G+C and GC3 content is observed in coronavirus genomes that results in low CAI values ^20,23^. Notably, SARS-CoV-2 and the other human coronaviruses also showed low tAIg and CSCg values that clearly distance themselves from human mRNAs, especially from highly expressed mRNAs such as ACTB (Figure 6A and Figure S6A). To analyze the effect of NSP1 on the expression of SARS-CoV-2 genes, we selected the mRNA encoding viral nucleocapsid (N) as representative since it is the most abundant viral mRNA detected in infected cells ^53^. The N mRNA includes the natural 80-nt 5’ leader (5’ L) that it is also present in the rest of viral mRNAs ^54^. Moreover, N mRNA shows a tAIg representative of full CDS of SARS-CoV-2 (≈ 21). As expected, expression of 5’ L-N was resistant to NSP1 WT (Figure 6B and C). Surprisingly, NSP1 M2 consistently reduced the expression of 5’ L-N to a similar extent to FFL, EGFP and RL reporters (Figures 1 and 6), confirming that expression of N mRNA is adapted to the presence of NSP1 WT. To test the contribution of 5’ UTR and codon usage on the resistance of N mRNA to NSP1, we replaced the 5’ L with 5’ UTR-82 alone or in combination with a variant of N that shows an improved codon usage for human cells (N1, tAIg= 0.248) (Figure 6B and C). Expression of 5’ UTR-82-N was now sensitive to NSP1 WT and M2 to a similar extent to FFL and RL reporters (Figures 6 and 1). Notably, the combination of 5’ UTR-82 and N1 dramatically increased the sensitivity to NSP1 WT (15- to 20-fold inhibition) comparable to that seen for EGFP and ACTB mRNAs (Figure 6B and C). We also found reduced levels of 5’ UTR-82-N1 mRNAs in the presence of NSP1 WT, showing that unlike parental 5’ L-N mRNA, 5’ UTR-82-N1 mRNA became sensitive to NSP1-induced degradation (Figure 6C). However, the replacement of 5’ UTR-82 with 5’ L significantly reduced the NSP1 sensitivity of the N1-expressing construct, showing that both natural 5’ L and codon usage of N mRNA contribute to resistance to NSP1.

**Figure 6.**
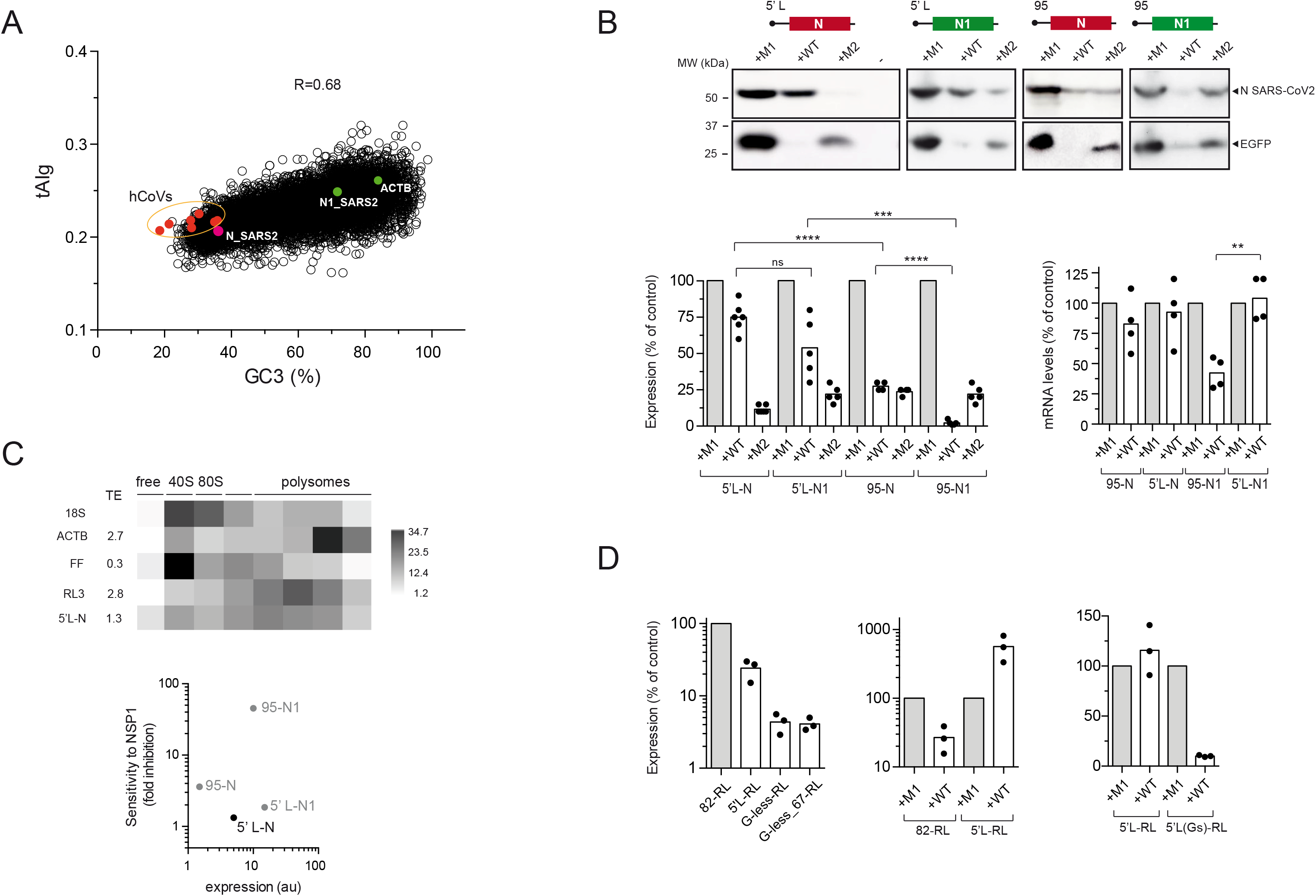
A combination of 5’ UTR and codon usage bias promotes resistance of N mRNA SARS-CoV-2 to NSP1. (A) Plot of tAIg and GC3 (%) distributions among human and coronavirus (hCoV) mRNAs. Values corresponding to human ACTB (green) and SARS-CoV-2 mRNA N (red) and its variant N1 (green) are indicated, along with the Pearson correlation. (B) A representative western blot analysis of N and EGFP accumulated in cells co-expressing NSP1 M1, WT or M2 (upper panel) and the corresponding quantification (bottom left panel). Relative quantification of mRNA levels of N variants in cells co-expressing NSP1 WT (bottom right panel). Significance was scored as described in previous figures. The combination of 5’ UTR (95 or 5’ L) with CDS N is indicated at the top of each panel. The tAIg values for N and N1 are 0.21 and 0.248, respectively. (C) Translation efficiency (TE) of 5’ L-N mRNA in HEK293T cells compared to some other reporters included in the co-transfection experiment (upper panel). Comparative analysis of the expression level and sensitivity to NSP1 among N variants with different combinations of 5’ UTR and codon usage. (D) Analysis of the expression of 5’ L-RL, G-less-RL and G-less_67-RL compared to 82-RL (left panel). The sensitivities of 5’ L-RL expression (middle panel) and its variant (5’ L(Gs)-RL, right panel) to NSP1 WT are shown.

Despite the suboptimal codon usage of natural N SARS-CoV-2 mRNA in human cells, polysome profiling experiments revealed a translation efficiency of N mRNA comparable to EGFP and much higher than FFL and RL mRNAs, which have similar tAIg (Figure 6C and supplementary table 1). To understand this, we first compared the expression of the four variants of N in HEK293T cells. The presence of 5’ L significantly increased the expression of the N protein to a level comparable to that found in 5’ UTR-82-N1. Despite sharing a similar nucleotide composition, 5’ L was more efficient that G-less /G-less_67 in directing the expression of RL (Figure 6D). Interestingly, NSP1 WT enhanced the expression of 5’ L-RL about 6-fold, reaching an expression similar to 5’ UTR-82-RL. However, when we placed a (GA)_6_ motif in the 5’ L-RL construct (5’L(Gs)-RL), a strong inhibition by NSP1 similar to the effect on G-less-M4-RL was observed. Moreover, we found an intermediate sensitivity of 5’ L to RocA compared to G-less and 5’ UTR 82 (Figure S7B). Taken together, these results suggest that SARS-CoV-2 mRNA has reached an optimal combination of 5’ UTR and codon usage bias to both support efficient translation and to resist the inhibitory effect of NSP1.

### A genome-scale model of NSP1 effect on translation and mRNA stability

To extend our analysis genome-wide, we used previously published datasets of mRNA abundance in cells expressing NSP1 alone ^31^ or in the context of infection ^27^ (Figure 7A). For this analysis, we selected only mRNAs whose 5’ UTR sequence was experimentally confirmed using nanocage technology (6000 mRNAs) ^55^. Notably, mRNAs with the greatest sensitivity to NSP1 (*down* group) showed shorter 5’ UTR and CDS than the median, and higher tAIg than the median in the two analyses (Figure S7A and B). Conversely, mRNAs whose levels increased upon SARS-CoV-2 WT infection or by NSP1 expression (*up* group) showed the opposite (Figure S7A and B). When comparing *down* and *up* groups, these differences were exacerbated (Figure S7A and B). Since tAIg of mRNAs influences ribosome elongation rates, we compared the sensitivity to NSP1 among mRNAs with predicted fast (short CDS and high tAIg) and slow (long CDS and low tAIg) ribosome transits. A dramatic difference in sensitivity to NSP1 was found between these two groups, especially in SARS-CoV-2-infected cells (Figure 7A), showing that mRNA with fast ribosome transits were particularly sensitive to NSP1-mediated inhibition. However, mRNAs with longer CDS and lower tAIg than the median showed resistance or even had increased levels in the presence of NSP1. The most abundant SARS-CoV-2 mRNAs fall into this group, showing a translation efficiency 2-3-fold higher in the presence of NSP1 ^27^.

**Figure 7.**
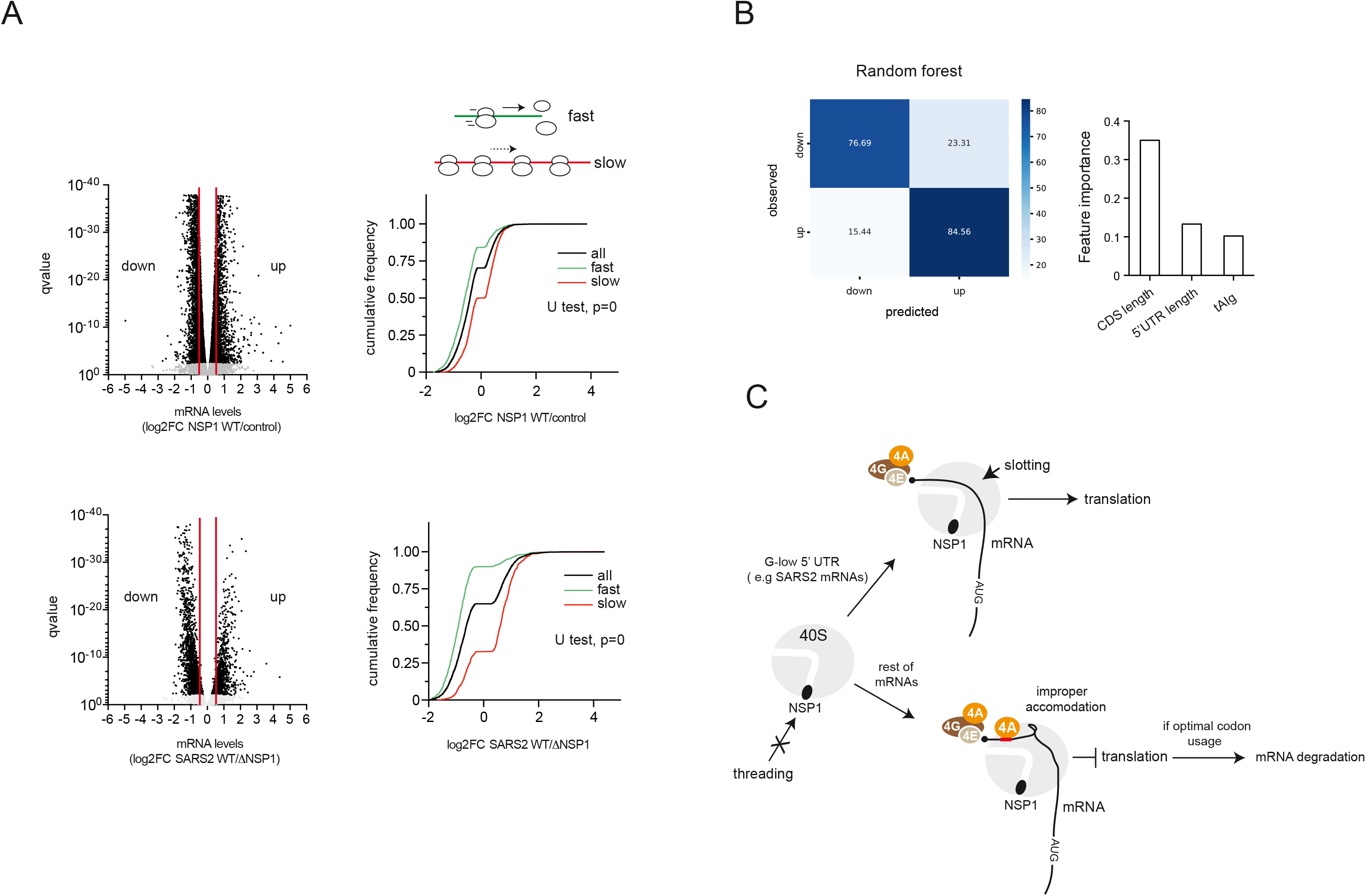
Genome-wide analysis of features in human mRNAs associated with sensitivity to NSP1. (A) Violin plots of the differential sensitivity of human mRNAs to NSP1 in transfected cells or in the context of infection. For the analysis, we used the datasets from Yuang *et al*. 2020 (top) and from Fisher *et al*. 2022 (GSE200422, bottom). mRNAs showing a log2FC <-0.5 or >0.5 were classified as *down* regulated or *up* regulated by NSP1, respectively. The cumulative fraction of log2FC values for mRNAs showing fast or slow ribosome transit for every dataset is shown. Mann-Whitney test for fast and slow groups is shown (B) Prediction of *down* and *up* classes by Random forest classifier. The mean accuracy obtained was 0.81. The resulting three most important features captured by the model are shown (C) Model of the differential effect of NSP1 on mRNA translation and stability. NSP1 binding prevents the penetration of mRNA through the entry channel of 40S (threading). However, lateral attachment of mRNA by slotting is still possible in the presence of NSP1. The resulting complex can proceed efficiently for mRNAs with G-low 5’ UTR. However, for mRNAs with eIF4A binding motifs (Gs or GAs, red) near the cap, slotting can result in an improper accommodation of the mRNA in the 40S groove that blocks translation initiation. When combined with optimal codon usage of the CDS, NSP1-mediated translation blockade can also result in mRNA degradation.

We also compared the nucleotide composition of 5’ UTR among *down* and *up* mRNA groups by computing the number of purines (G alone or G+A) throughout the 5’ UTR using a sliding 5-nt window to detect mRNAs enriched in eIF4A-binding motifs. Remarkably, the mRNA *down* group showed a significant enrichment in G content from +11 to +18 position with respect to the mRNA 5’ cap (Figure S7A and B). This finding is in good agreement with what described in Figure 5 using G-less-RL variants with Gs and GA motifs near the 5’ cap.

Finally, we used supervised machine learning models to predict the sensitivity of human mRNAs to NSP1 based on the discovered features. Among all models tested, Random Forest classification generated the best performance using the dataset of Fisher *et al*. 2022. The model predicted *down* and *up* classes with a high accuracy (81%) by selecting CDS length, tAIg and 5’ UTR length as features with the greatest importance in the classification process (Figure 7B). The accuracy increased to 83% when features were transformed using PCA (principal component analysis) before training the classifier. Taken together, these analyses and predictions support the model depicted in Figure 7.

## Discussion

Among all known viral proteins that interfere with host translation, SARS-CoV-2 NSP1 is unique in the way it blocks cellular translation initiation, which may also result in mRNA degradation ^56^. Taken together, our results show that the effect of NSP1 on translation initiation depends on the length and nucleotide composition of the 5’ UTR, reflecting a differential mode of 43S-PIC recruitment to mRNAs, whereas NSP1-mediated mRNA degradation is strongly influenced by codon usage bias. We propose the existence of both threading and slotting mechanisms of 43S-PIC recruitment to mRNA that are differentially affected by NSP1. According to our model, the presence of eIF4A-binding G/GA-motif near the 5’ cap promotes (or enforces) the threading of mRNA through the entry channel during 43S-PIC recruitment. When the mRNA entry channel of 43S-PIC is blocked by NSP1, eIF4A-bound mRNA could not be properly accommodated within the 40S groove, resulting in an aberrant 48S-PIC that may eventually undergo mRNA degradation (Figure 7C); some previous data support this model. First, NSP1 binding to 40S did not prevent mRNA attachment so that the eventual mRNA degradation occurred on the 48S-PIC ^36,57,58^. Second, eIF4A has been detected near the mRNA entry channel in 48S-PIC assembled with alphavirus or synthetic mRNAs ^42,59^. Third, the presence of eIF4A as part of the eIF4F complex increased the sensitivity to NSP1 *in vitro* even for those mRNAs that do not require eIF4F for translation ^36^.

When combined with a threading-prone 5’ UTR, optimal codon usage exacerbated NSP1-induced mRNA degradation. In our opinion, the simplest explanation for this is the positive influence that fast elongating CDS could have on translation initiation, feeding mRNA with new ribosomal subunits and eIFs for reinitiation that amplifies the inhibitory signal of NSP1 on target mRNAs. This idea is further supported by the finding that mRNAs with predicted fast ribosome transit showed a dramatic sensitivity to NSP1-induced mRNA degradation. However, it is important to note that optimal codon usage did not promote NSP1-mediated mRNA degradation when combined with a slotting-prone 5’ UTR. This shows that susceptibility of a given mRNA to NSP1-induced degradation critically depends on the way 43S-PIC is recruited to the mRNA. Although our data do not reveal the mechanistic details of NSP1-induced mRNA degradation, the fact that NSP1 can also promote stabilization of some mRNAs (e.g., SL50-FFL and G-less-RL) suggests that beyond an intrinsic activity of NSP1 bound to 40S, mRNA degradation could be the result of a safeguard activity associated with 48S-PIC to remove improperly accommodated mRNA within the 40S groove. This activity would require eIF4A (and perhaps DDX3) bound to mRNA and the participation of eIF3G as described recently ^36^.

According to structural model of 40S-NSP1 complex, lateral accommodation of mRNA by slotting is sterically possible when mRNA entry channel is blocked by NSP1 ^29,30^. Our data strongly suggest that NSP1 allowed or even stimulated the slotting of 43S-PIC on G-less/low 5’ UTR mRNAs, including those of SARS-CoV-2 bearing the 5’ L. Despite they are unrelated in sequence, the striking compositional and functional similarities between G-less/low 5’ UTR and viral 5’ L are suggestive of an evolutionary adaptation of SARS-CoV-2 to exploit the ability of 43S-PIC to be slotted on certain mRNAs. Our results support the notion that, more than a specific participation of the 5’-proximal stem loop (SL1) in the translational resistance of 5’ leader mRNA to NSP1 ^25,40^, the slotting-prone activity of coronaviral 5’ L mRNA together with a suboptimal codon usage of CDS would provide an effective mechanism to escape the effect of NSP1. However, unlike reporter genes with G-less/low 5’ UTR that are translated less efficiently, 5’ L of SARS-CoV-2 directed efficient translation to reporter mRNA. We propose that the presence of G-poor secondary structure (SLs) of moderate stability in the 5’ L of SARS-CoV-2 mRNAs provides an optimal combination of translation efficiency and resistance to NSP1. Further studies on these adaptive trends could be useful to better understand long-term evolutionary trajectories of Coronavirus and other viruses infecting humans that show similar trends.

## Acknowledgments

We thank Fernando Rodríguez Pascual and Araceli del Arco (both at CBM, Madrid) for providing us with pNL1.1.CMV plasmid and goat anti-luciferase antibody, respectively. We also thank Irene Díaz-López for suggestions and sharing ideas.

## Author contributions

I.V. conceived the study, directed the research, performed most of the experiments and wrote the manuscript. J.B. performed most of the western blots. T.M. performed many mRNA quantifications and polysome analysis. R.T. wrote and executed scripts for bioinformatic analysis. M.R.-P. and M.S. designed and performed infections with SARS-COV-2. M.N.-B. performed machine learning simulations.

## Materials and Methods

### Plasmids and recombinant DNA

Plasmids encoding SARS-CoV-2 NSP1 variants were generated by exchanging the EGFP coding sequence of pEGFP-N1 plasmid (Clontech) for the corresponding nsp1 genes. For this, nsp1 coding sequences were amplified by PCR from pDONR207 SARS-CoV-2 nsp1 (WT, Addgene #141255), pDONR207 SARS-CoV-2 nsp1 Delta RC (M1, K164A/H165A, Addgene #164522) and pDONR207 SARS-CoV-2 nsp1 R124A/K125A (M2, Addgene #164523) plasmids using the primers 5-EX-NSP1 and 3-NB-NSP1 and cloned into pEGFP-N1 digested with *EcoR*I and *Not*I enzymes. Some of the reporter plasmids used here have been described previously: pFFL(firefly luc), pRL (renilla luc), p242-RL, p427-RL, p657-RL, pG-less-RL, pG-low-FFL, pSt20-RL/FFL and pSt30-Rl/FFL ^14^. pEGFP-N1 and pDsRed-N1 were purchased from Clontech and pbeta-Gal expresses the lacZ gene under the CMV-IE promoter. The new reporter plasmids generated in this study were:

-pNanoLuc. The coding sequence of NanoLuc was amplified by PCR using the primers 5-NanoL and 3-NanoL from pNL1.1.CMV plasmid (Promega). The resulting fragment was digested with *BamH*I and *Not*I enzymes and cloned into pEGFP-N1 digested with the same enzymes, resulting in the replacement of EGFP for NanoLuc.

-pACTB. Human ACTB CDS was amplified by RT-PCR using the primers 5-N-hACTB and 3-X-hACTB and cloned into a plasmid bearing a myc epitope using *Not* I and *Xba* I enzymes. Then, the *BamH*I-*Not*I fragment was subcloned into pEGFP-N1 digested with the same enzymes. The resulting plasmid encodes a myc-tagged version of ACTB.

-pN SARS-2. The coding sequence of N gene with the natural codon usage of SARS-CoV-2 from pSARS-CoV-2 (N) plasmid (Addgene #153201) was digested with *BamH*I and *Not*I enzymes and cloned into pEGFP-N1 using the same enzymes.

- pG-low-RL/EGFP. The *Nde*I-*BamH*I fragment from pG-low-FFL plasmid was subcloned into pRL and pEGFP-N1 plasmids digested with the same enzymes.

All variants of pG-less-RL, p5’ L-N, p5’L-RL, p5’L(Gs)-RL, TISU-EGFP, Slot-RL and the recoded versions of pEGFP (pEGFP-1, EGFP-2), pRL (pRL1, pRL2 and pRL3) and N (p5’L-N1, pN1) were obtained from GenScript.

-pTISU-RL. A PCR fragment including TISU element from pTISU-EGFP plasmid was cloned into pRL using *Nde*I and *Sac*I enzymes.

All constructs generated in this work were verified by sequencing.

### Cell culture and transfection

HEK293T, HeLa, MEFs, Calu-3 and Vero E6 cells were grown in DMEM supplemented with 10% fetal calf serum and antibiotics. Cells were authenticated by microscopic examination and the absence of mycoplasma contamination was confirmed every two months of culture. For plasmid transfection, cells growing in a P24 plate were transfected with 1 µg of plasmid (or a combination of plasmids) and 2 µL of Turbofect (HEK293, MEFs and HeLa) or 2 µL of Lipofectamine 2000 (Calu-3 and VeroE6) according to the manufacturer’s recommendation (ThermoFisher and Invitrogen, respectively). For co-transfections, we combined up to 0.5 µg of pNSP1 plasmids with a mix of up to three reporter plasmids (0.15-0.2 µg each). We always included pEGFP-N1 plasmid as a readout of transfection efficiency and sensitivity to NSP1. For titration with increasing amounts of pNSP1-WT or M2, the transfection mixtures were filled in with pNSP1-M1 (null) plasmid up to 0.5 µg. Transfected cells were incubated 24 h at 37 °C and extracts were prepared for reporter activity measurement, western blot or RNA extraction. Oligonucleotide transfection was carried out as described before ^14^.

### SARS-CoV-2 infection

Vero E6 cells growing in P24 wells were infected with SARS-CoV-2 (Wuhan-1) at a moi = 2 pfu/cell in 0.5 mL OPTIMEM (Gibco) under biosafety level 3 conditions. After 1 h adsortion, the inoculum was removed and cells were transfected with a combination of pEGFP, pRL3- and pFFL plasmids (0.25 μg each) using Lipofectamine 2000 (Invitrogen). Cell extracts were prepared at 16 or 24 hpi for luciferase activity measurement and western blot.

### Measurement of reporter gene expression

Enzymatic measurements were carried out in cell extracts prepared in passive lysis buffer (Biotium). Firefly, renilla and NanoLuc luciferases were measured in a Berthold luminometer using D-luciferin (Promega), collantrazine (Biotium) and Nano-Glo (Promega), respectively. To avoid luminometer saturation, extracts were diluted to keep luciferase activities below 2x10^6^ RLUs. Beta-galactosidase activity was measured using ONPG as substrate (Promega). EGFP and dsRed accumulation was measured by western blot (see below) and by fluorescence emission of extracts using a GloMax microplate reader (Promega). In both cases, a standard curve was included for relative quantification in the linear range of measurement.

### Western blot

Western blots were carried out as described previously (Díaz-López et al., 2019) using the following primary antibodies: anti-EGFP (11814460001, Roche), anti-dsRed (a gift from José María Requena, CBMSO, Madrid.), anti-FFLuc (a gift from Araceli del Arco, CBMSO, Madrid), anti-N SARS-CoV-2 (MA5-38034, Invitrogen), anti-NSP1 SARS-CoV-2 (STJ11103222, St. John’s lab), anti-myc (05-724MG, Merck-Millipore), anti-eIF4A (STJ2724, St. John’s lab,) and anti-eIF3G (STJ23512, St. John’s lab) and anti-eIF2α (sc-11386, Santa Cruz Biotechnology). Blots were developed with Luminata™/Immobilon® Crescendo Western HRP Substrate (Millipore) and bands were quantified by densitometry using the linear range of the signal.

### RNA extraction and quantitative RT-PCR

Total RNA was extracted from cells growing in P24 or P12 wells using the RNeasy plus kit (QIAGEN). The resulting RNA was digested with 2 U of DNAse I for 30 min at 37 °C and further purified using RNA clean & concentrator 5 columns (Zymo research). About 1.5 μg of total RNA was retrotranscribed with hexamers using Superscript VI retrotranscriptase (Invitrogen). qPCR amplification was carried out with GoTaq qPCR mix (Promega) in a BioRad CFX-384 thermocycler using a 1/200 dilution of corresponding cDNAs. Relative quantification was performed using the Relative Expression Software Tool (REST) ^60^ and the following formula:

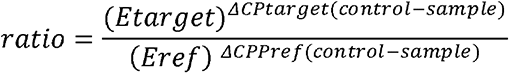

18S RNA was used as reference gene. For experiments analyzing the effect of NSP1 WT on reporter mRNA accumulation, samples expressing NSP1 M1 (null mutant) were used as control.

### Measurement of translation efficiency of mRNAs

Translation efficiencies (TE) of mRNAs were calculated by polysome profiling of HEK293T extracts as described previously ^14^. Briefly, one P100 plate of HEK293T cells were transfected with a combination of up to 4 different reporter expressing plasmids for 24 h. Then, monolayers were lysed in polysome buffer (Tris-HCl 30 mM pH 7.5, 100 mM KCl, 5 mM MgCl_2_, 1 mM DTT, 1% Triton X-100 and 50 μg/ml cycloheximide) for 15 min on ice. Cell lysates were clarified by low-speed centrifugation and spun on a 10-40% sucrose gradient at 35,000 rpm for 3 h at 4 °C in an SW40 rotor. Ten fractions were collected using an ISCO fractionator coupled to a UV recorder, followed by trizol extraction and ethanol precipitation. The resulting RNA pellets were resuspended in water and digested with 2 U of DNAse I for 30 min at 37 °C followed by purification using RNA clean & concentrator 5 columns (Zymo Research). mRNA levels in every fraction was quantified by RT-qPCR as described above. TE for every mRNA was calculated as the ratio between the relative amount found in polysomal region (fractions from 7 to 10) and monosomal (80S) and submonosomal regions (fractions from 2 to 5).

### Codon usage, tRNA adaptation index (tAI) and codon stability coefficient (CSC) calculations

Codon usage tables of human mRNAs were downloaded from HIVE-CUT repository ^61^. Coronavirus coding sequences were extracted from reference genomes deposited in GenBank (SARS-CoV-2:NC_045512.2., SARS-CoV1: NC_004718., MERS: NC_019843.3., HCoV-OC43: NC_006213., HCoV-229E: NC_002645., HCoV-NL63: NC_005831., HCoV-HKU1: NC_006577). Nucleotide composition, codon usage per thousand and codon adaptive index (CAI) of coronavirus genomes and reporter genes were calculated using the tools hosted in the CAIcal server ^62^. Transcript-level average tAI (tAIg) and CSC (CSCg) were calculated using the following formula:

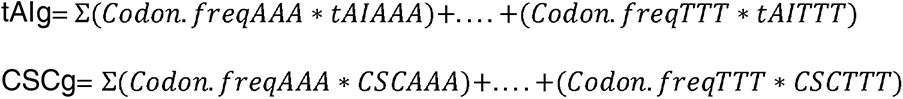

We used tAI and CSC values for all 61 non-stop codons previously calculated using experimental measurements of tRNA levels and mRNA half-lives in HEK293T cells, respectively ^43,44^.

### Recoding of reporter genes

Codon usage optimization and deoptimization of EGFP, RL and N SARS-CoV-2 coding sequences were carried out using the harmonization tool described in http://www.codons.org/ ^63^. We constructed a codon usage table of our HEK293T cells that included all mRNAs whose expression was detected by RNAseq.^14^. This table was used as reference for the recoding process.

### Data processing and bioinformatic analysis

We used raw and processed datasets deposited in GEOs. Data normalization per kilobase per million reads (RPKM) was applied where necessary. Differential expression was calculated using DESeq2 and expressed as log2FC ^64^. For further analysis, we selected only mRNA whose 5’ UTR sequence was experimentally confirmed using nanocage technology (5000-6000 mRNAs depending on the dataset) ^55^. 5’ UTR features including length and nucleotide composition were calculated using excel formulas. A Python script was prepared to compute the number of G and G+A in 5’ UTR datasets using a 5 nt-sliding window. RNA secondary structure was predicted using the RNAfold program (ViennaRNA package). Human mRNAs with predicted fast ribosome transit were defined as those that showed a CDS length below the lower limit of the 95% CI of the median and a tAIg above the upper limit of the 95% of the median. mRNAs with predicted slow ribosome transit were defined as those that showed a CDS length above the upper limit of the 95% CI of the median and a tAIg below the lower limit of the 95% CI of the median.

### Machine learning models

A machine learning pipeline was constructed in Python using scikit-learn ^65^. As output variable, we selected data deposited in GSE200422 corresponding to the effect of SARS-CoV-2 WT *vs* ΔNSP1 infection on mRNA abundance in human Calu-3 cells at 8 hpi ^27^. The dataset was filtered for a q-value <0.01. We incorporated 5 features of 5’ UTR and 5 features of CDS of every mRNA: 5’ UTR length (nt), 5’ UTR G+C (%), predicted secondary structure of 5’ UTR (ΔG°), number of Gs in 5-25 of 5’ UTR, number of G+A in 5-25 of 5’ UTR, CDS length (number of codons), CDS G+C (%), CDS GC3 (%), tAIg_HEK293 and CSCg_HEK293. For Random Forest classification, each feature was standardized so that the values were scaled to a mean of 0 and a standard deviation of 1. Continuous values (Log2FC) of output variable were transformed into two categorical classes: down (=0), showing log2FC < -0.5 and up (=1) showing log2FC >0.5. The two classes were then balanced resulting in a dataset of 1652 genes. Then, the balanced dataset was split into training and testing sets using a ratio of 2.1 (two-thirds training and one-third testing). The hyperparameters for the Random Forest classifier were optimized using a grid search with 10-fold cross-validation. To determine feature importance and their contribution to the model, the mean decrease Gini index was used. CDS length was recognized as the most important feature, followed by 5’ UTR length and tAIg.

### Statistical analysis

Statistical analyses were carried out in GraphPad Prism 6. Two-tailed *t*-test and non-parametric tests were performed using at least 3 biological replicates per group and a least two technical replicates per sample. Significance was scored as follow: p<0.0001, ***: p<0.001, **: p< 0.01, *: p<0.05, ns = not significant. Pearson and Spearman correlations were calculated using default parameters.

---

**Figure S1.**
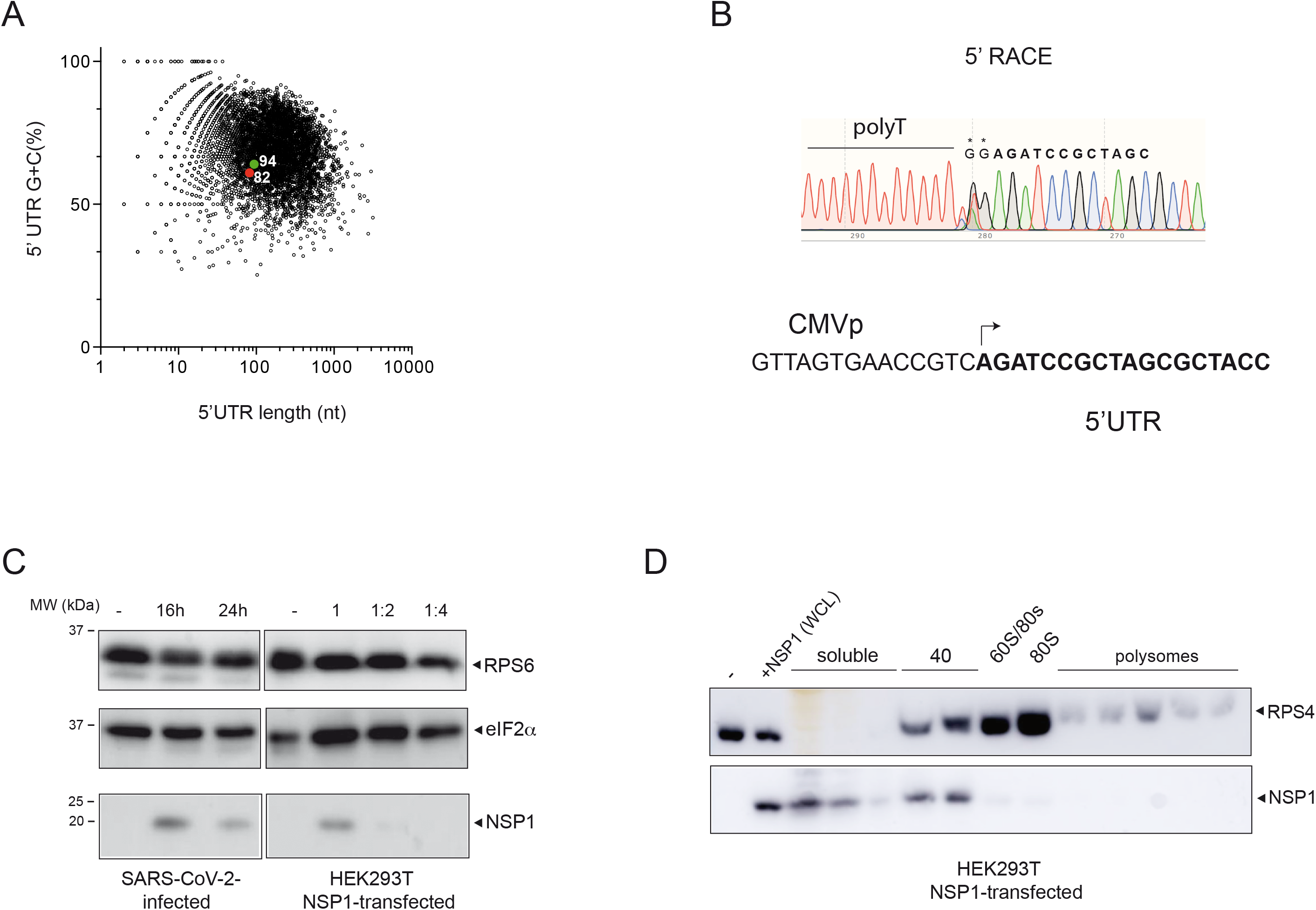
(A) Plot of length and G+C composition of human 5’ UTRs previously identified by Nanocage technology ^55^ that were used in this study. The positions of parental 94- and 82-5’ UTR of reporter mRNAs are shown. (B) Mapping the 5’ end of 82-RL mRNA by 5’ RACE. The sequencing chromatogram is shown. Asterisks denote the extra GG that results from known template-independent addition of two Cs by reverse transcriptase during cDNA synthesis ^66^. The mapped transcription start site (arrow) matched with that described for parental pEGFP-N1(Clontech). (C) Comparative analysis of NSP1 levels accumulated in HEK293T cells transfected with pNSP1-WT plasmid and VeroE6 infected with SARS-CoV-2. The 1/2 and 1/4 dilutions of HEK293T extracts are indicated. Blots were probed with the indicated antibodies. (D) Association of NSP1 to 40S subunits in transfected HEK293T cells. Fractions from polysome analysis were probed with anti-NSP1 and anti-RPS4 antibodies. The positions of the resulting identified peaks are shown.

**Figure S2.**
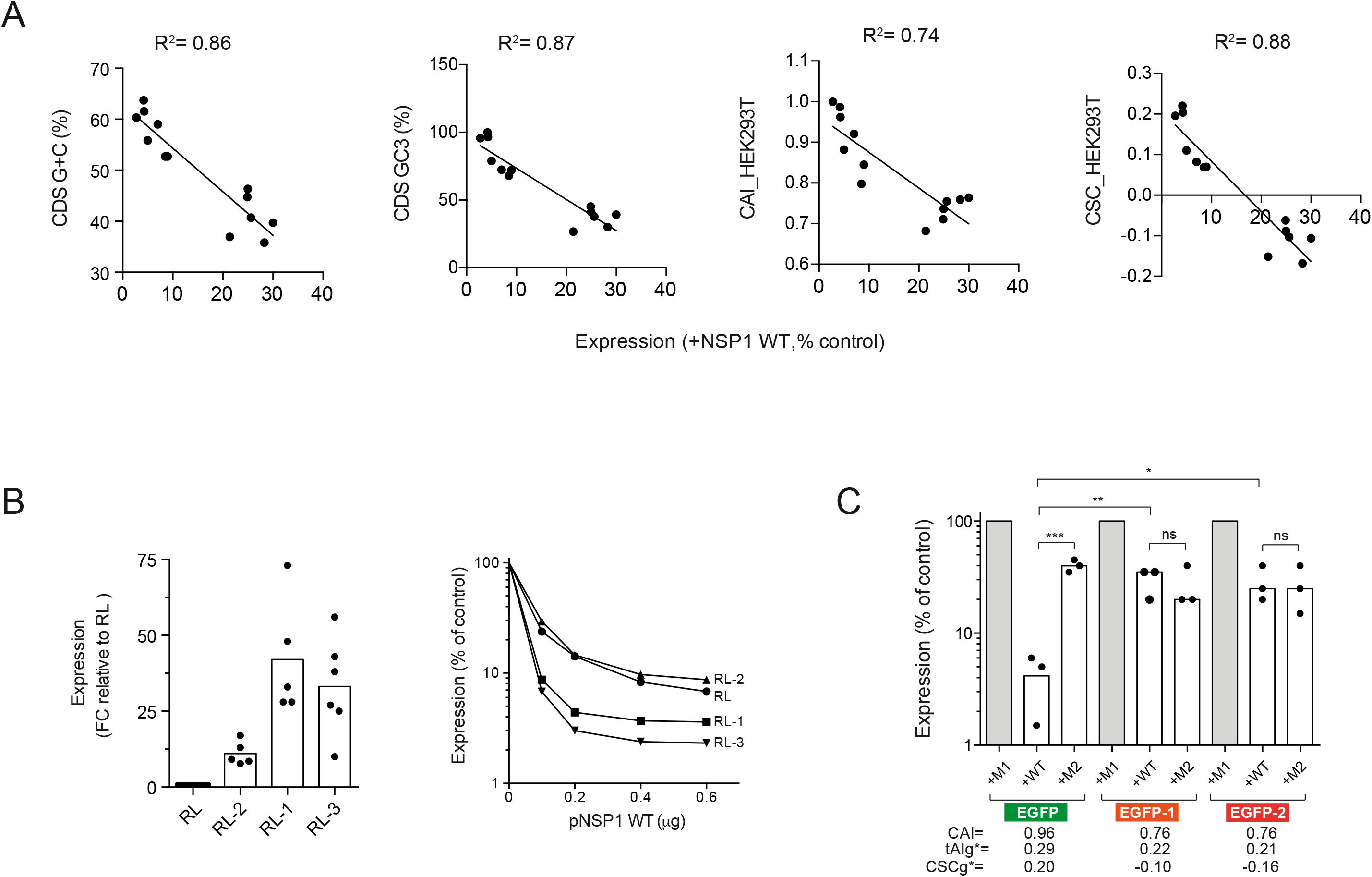
(A) Correlation between sensitivity to NSP1-WT and reporter’s CDS features including G+C (%), GC3 (%), CAI and CSCg estimated for HEK293T cells. The resulting Pearson R^2^ is shown. (B) Expression of RL variants (RL-1 to RL-3) with different codon usage bias. Data are expressed as fold change (FC) relative to RL expression (left panel). Effect of increasing amount of pNSP1 WT plasmid on expression of RL variants (right panel). (C) Effect of EGFP CDS recoding on sensitivity to NSP1 WT and M2. The CAI, tAIg and CSCg values for parental (EGFP, green) and EGFP-1 and EGFP-2 (orange to red) are indicated. Significance was scored as described in Materials and Methods.

**Figure S3.**
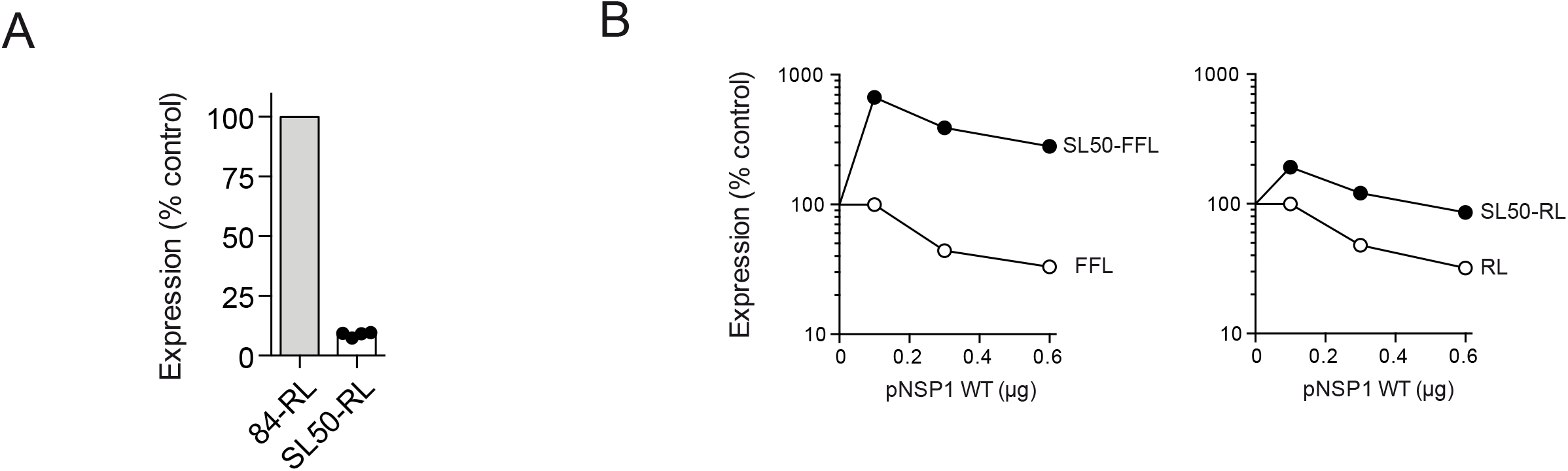
(A) Effect of upstream SL50 on RL expression. Data are expressed relative to parental 82-RL (100%). (B) Comparative analysis of the effect of increasing amounts of pNSP1 WT plasmid on expression of SL50-FFL and SL50-RL relative to pNSP1 M1(100%).

**Figure S4.**
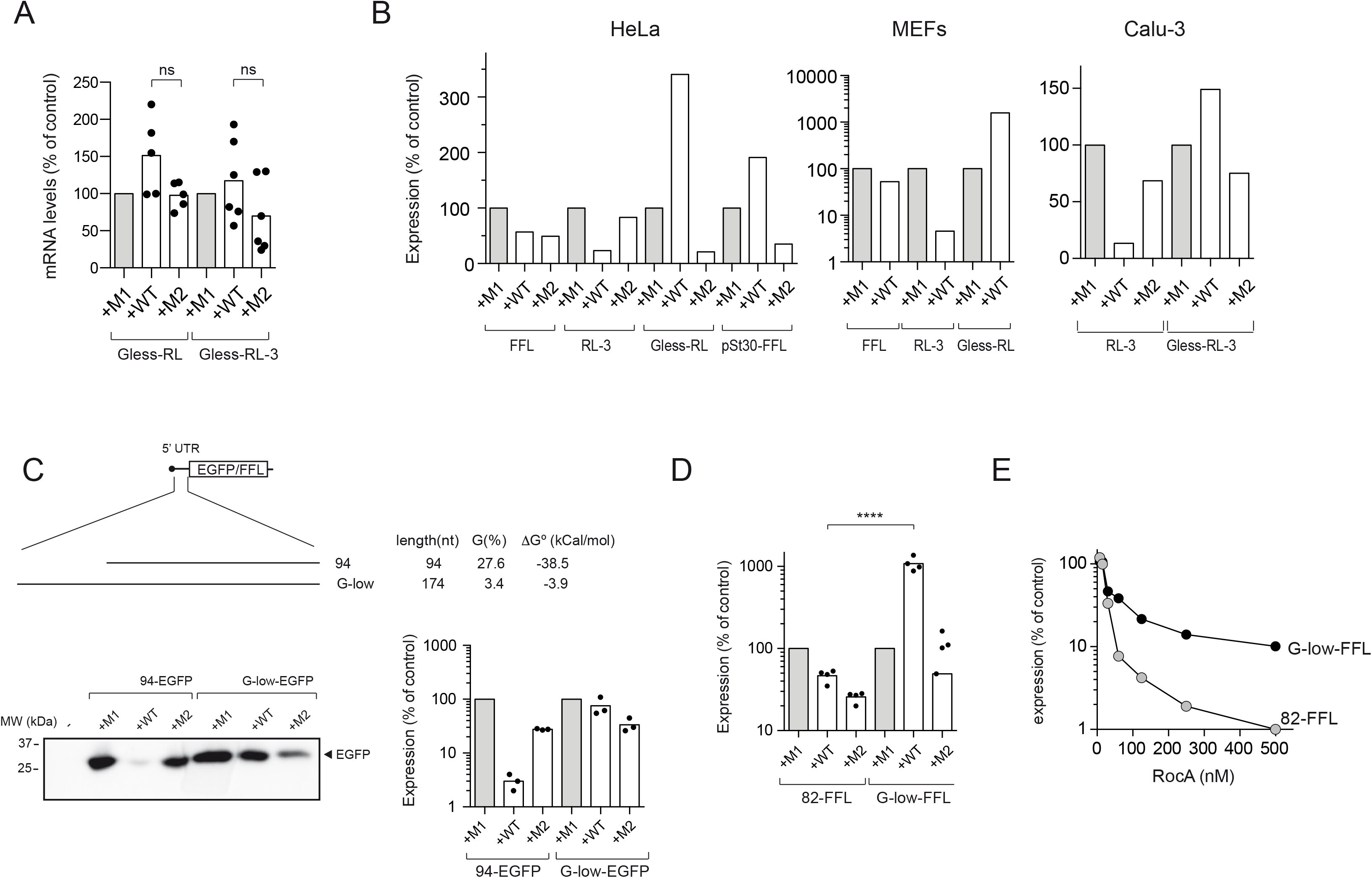
(A) Relative mRNA levels of G-less-RL and G-less-RL3 accumulated in cells co-expressing NSP1 WT or M2. Data were expressed and significance was scored as described in Figure 1. (B) Effect of NSP1 WT and M2 on expression of the indicated reporters in HeLa, MEF and Calu-3 cells. Data are the mean from two (HeLa and MEFs) or three (Calu-3) independent experiments. (C) Effect of G-low 5’ UTR on the sensitivity of EGFP and FFL to NSP1 WT and M2. Features of 94 and G-less 5’ UTRs including length, number of Gs and stability (ΔG° in kcal/mol) are indicated (upper panel). A representative western blot of EGFP accumulated in cells co-transfected with the indicated constructs and the resulting quantification from three independent experiments are shown (bottom panels). (D) Influence of G-low 5’ UTR on the sensitivity of FFL to NSP1 WT and M2. Data are expressed and significance was scored as described in previous figures. (E) Comparative analysis of the sensitivity of 82-FFL and G-low-FFL to RocA. Inhibitor addition and measurements were taken as described above.

**Figure S6.**
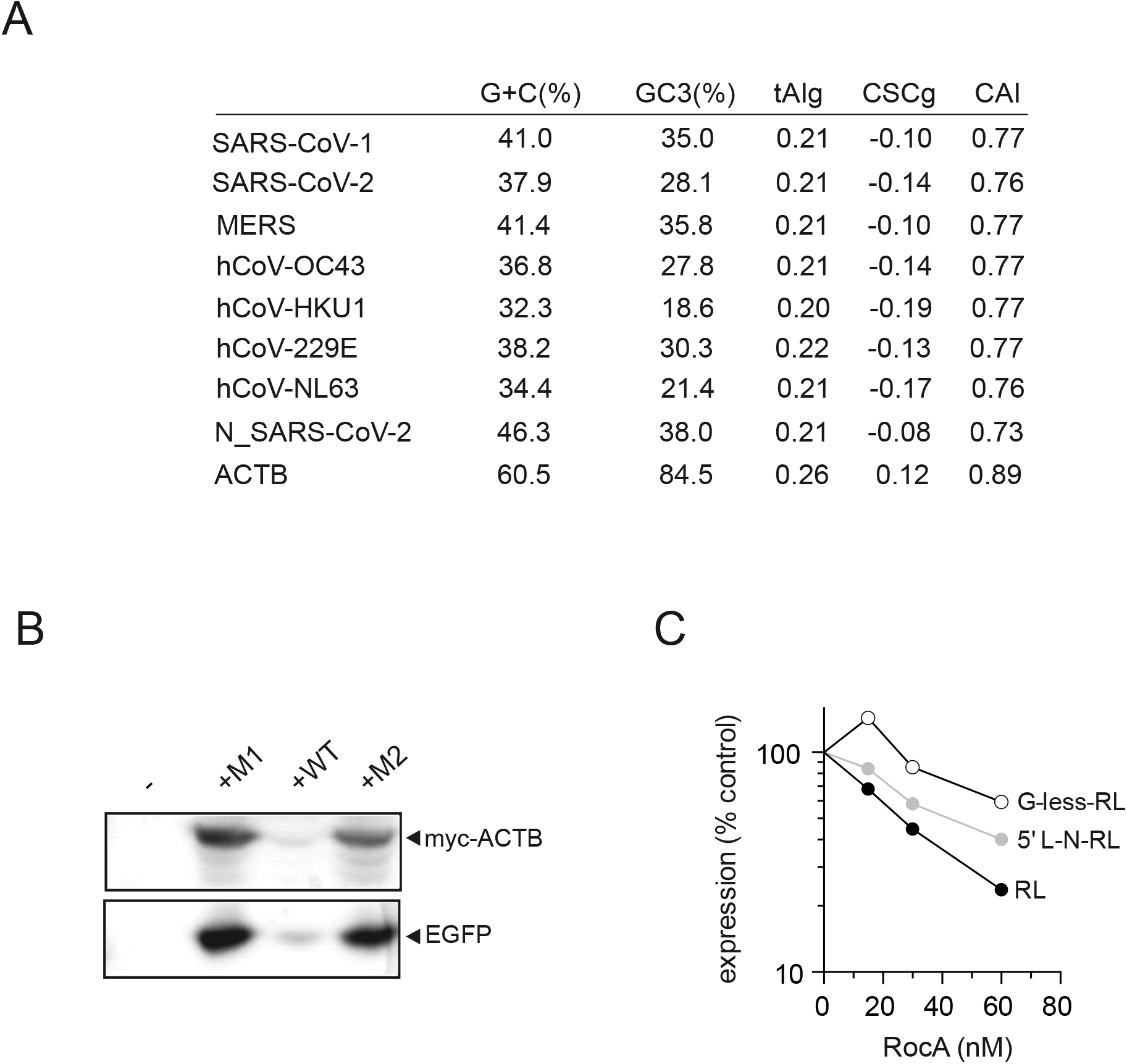
(A) Nucleotide composition and codon usage metrics of coding sequences of representative human coronavirus and ACTB mRNAs. (B) Sensitivity of human ACTB to NSP1 and its mutants. A representative western blot of myc-ACTB and EGFP is shown. (C) Comparative analysis of the differential sensitivity of RL, G-less-RL and 5’ L-RL expression to RocA.

**Figure S7.**
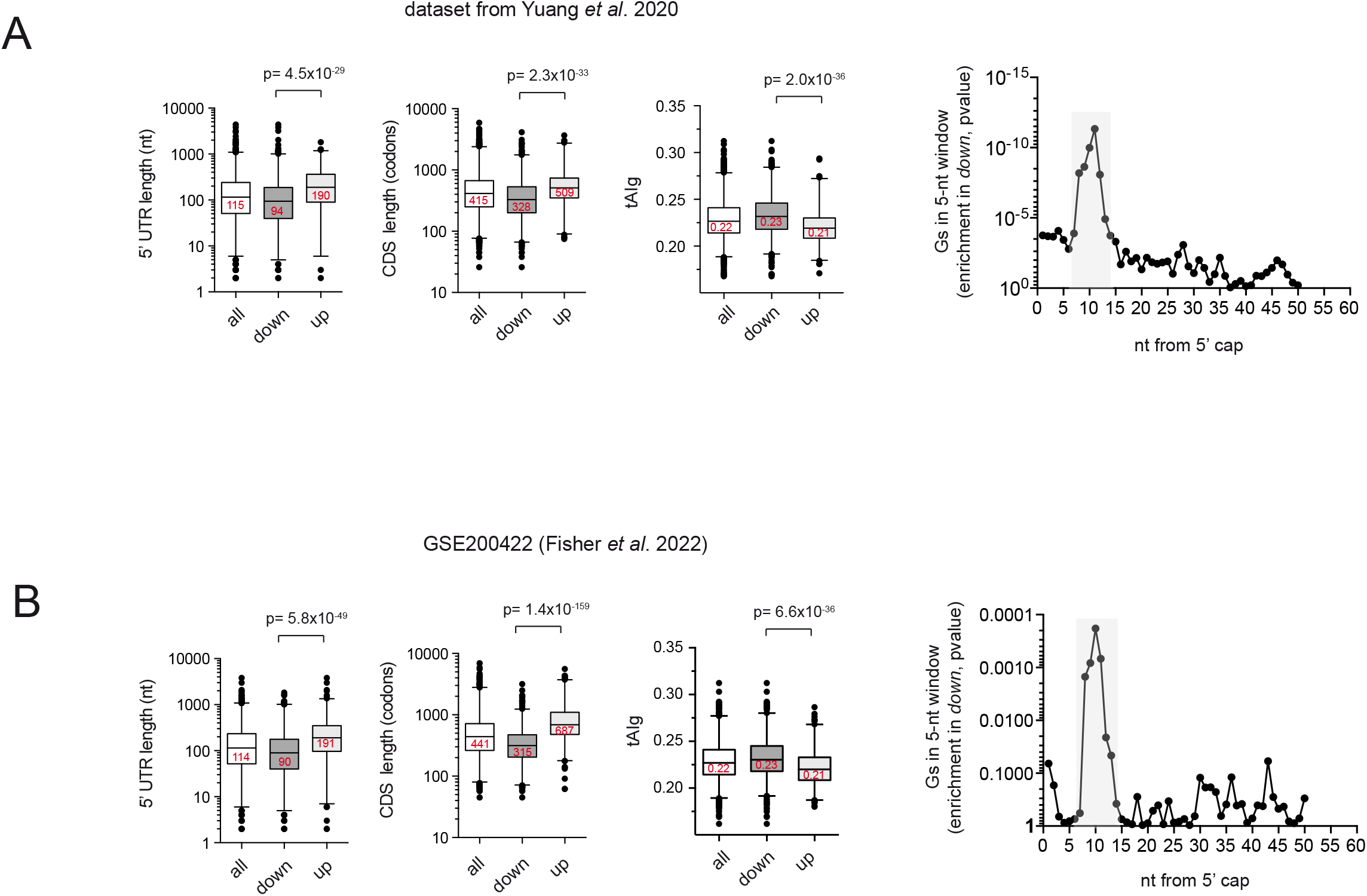
Comparative analysis of 5’ UTR length, CDS length (in codons) and tAIg in all, *down* and *up* groups using the indicated datasets of mRNA abundance. Plot boxes represent the median and the 1-99 percentile whiskers; p-values after Mann-Whitney U test are indicated. The enrichment in Gs along the first 50 nt of the 5’ UTR in the *down* group of mRNA for every analysis is also shown. Data represent the p-value after a *t*-test comparison between *down* and *up* groups.

