## Supplementary table 1 for "The differential effect of SARS-COV-2 NSP1 on mRNA translation and stability reveals new insights linking ribosome recruitment, codon usage and virus evolution"

| reporter | coding sequence | G+C(%) | tAIg | CAI | CSCg |
|---|---|---|---|---|---|
| EGFP(parental) | ATGGTGAGCAAGGGCGAGGAGCTGTTCACCGGGGTGGTGCCCATCCTGGTCGAGCTGGACGGCGACGTAAAC<br>GGCCACAAGTTCAGCGTGTCCGGCGAGGGCGAGGGCGATGCCACCTACGGCAAGCTGACCCTGAAGTTCATC<br>TGCACCACCGGCAAGCTGCCCGTGCCCTGGCCCACCCTCGTGACCACCCTGACCTACGGCGTGCAGTGCTTC<br>AGCCGCTACCCCGACCACATGAAGCAGCACGACTTCTTCAAGTCCGCCATGCCCGAAGGCTACGTCCAGGAG<br>CGCACCATCTTCTTCAAGGACGACGGCAACTACAAGACCCGCGCCGAGGTGAAGTTCGAGGGCGACACCCTG<br>GTGAACCGCATCGAGCTGAAGGGCATCGACTTCAAGGAGGACGGCAACATCCTGGGGCACAAGCTGGAGTAC<br>AACTACAACAGCCACAACGTCTATATCATGGCCGACAAGCAGAAGAACGGCATCAAGGTGAACTTCAAGATC<br>CGCCACAACATCGAGGACGGCAGCGTGCAGCTCGCCGACCACTACCAGCAGAACACCCCCATCGGCGACGGC<br>CCCGTGCTGCTGCCCGACAACCACTACCTGAGCACCCAGTCCGCCCTGAGCAAAGACCCCAACGAGAAGCGC<br>GATCACATGGTCCTGCTGGAGTTCGTGACCGCCGCCGGGATCACTCTCGGCATGGACGAGCTGTACAAGTAA | 61.53 | 0.29 | 0.96 | 0.20 |
| EGFP-1 | ATGGTGTCAAAAGGAGAAGAGCTGTTCACGGGAGTTGTACCAATATTAGTGGAATTAGACGGCGATGTTAAC<br>GGACATAAATTCTCAGTTTCCGGGGAAGGAGAAGGGGATGCCACATACGGTAAATTGACACTTAAATTTATT<br>TGTACAACAGGAAAATTGCCTGTTCCTTGGCCTACATTGGTGACAACATTGACATACGGAGTTCAATGTTTT<br>TCTAGATACCCTGATCACATGAAACAACACGATTTCTTCAAATCGGCCATGCCTGAAGGATACGTCCAGGAA<br>CGCACCATTTTCTTCAAGGATGATGGTAACTACAAAACAAGAGCCGAAGTGAAATTTGAGGGGGATACTTTA<br>GTTAATAGAATCGAATTGAAAGGAATAGACTTCAAAGAGGATGGCAACATTTTGGGACATAAGCTTGAATAC<br>AACTACAATTCCCATAACGTGTATATCATGGCGGATAAACAAAAGAACGGGATTAAGGTAAATTTTAAGATT<br>AGACACAACATAGAAGATGGATCCGTGCAACTGGCCGATCATTACCAACAAAACACACCTATTGGAGATGGA<br>CCTGTTTTGTTACCTGATAACCATTACTTGTCTACACAATCTGCGTTGTCTAAGGATCCTAACGAAAAAAGA<br>GACCATATGGTTCTGTTAGAATTTGTGACAGCAGCTGGCATTACACTAGGTATGGATGAGCTCTACAAATAA | 39.72 | 0.22 | 0.76 | -0.10 |
| EGFP-2 | ATGGTTAGTAAAGGTGAGGAACTCTTTACCGGGGTTGTTCCTATTTTGGTTGAATTGGATGGAGACGTCAAC<br>GGACATAAATTTTCTGTCTCTGGAGAAGGAGAAGGAGATGCCACATACGGAAAATTGACATTGAAATTTATT<br>TGTACCACTGGAAAATTGCCTGTTCCATGGCCAACCTTGGTTACAACATTAACATACGGAGTTCAATGTTTT<br>TCTAGATACCCAGATCATATGAAACAACATGATTTTTTTAAAAGTGCGATGCCTGAAGGGTACGTGCAGGAA<br>AGAACAATTTTTTTTAAAGATGACGGAAACTACAAAACAAGAGCCGAAGTTAAATTTGAAGGAGATACATTG<br>GTTAATAGAATTGAATTGAAAGGAATTGATTTTAAAGAAGATGGAAACATTTTAGGACACAAATTGGAATAC<br>AACTACAACTCTCACAATGTTTACATTATGGCAGACAAACAAAAAAACGGAATTAAAGTTAACTTTAAAATT<br>AGACATAACATTGAAGATGGATCCGTGCAATTGGCCGATCATTACCAACAAAATACACCTATTGGAGATGGA<br>CCTGTTTTGTTGCCTGATAACCATTACTTGTCCACACAATCTGCCTTGTCTAAAGACCCGAACGAAAAAAGA<br>GATCATATGGTTTTACTTGAATTTGTTACAGCGGCGGGAATTACTTTGGGCATGGACGAATTATATAAATAA | 35.83 | 0.21 | 0.76 | -0.16 |
| FFL | ATGGAAGACGCCAAAAACATAAAGAAAGGCCCGGCGCCATTCTATCCTCTAGAGGATGGAACCGCTGGAGAG<br>CAACTGCATAAGGCTATGAAGAGATACGCCCTGGTTCCTGGAACAATTGCTTTTACAGATGCACATATCGAG<br>GTGAACATCACGTACGCGGAATACTTCGAAATGTCCGTTCGGTTGGCAGAAGCTATGAAACGATATGGGCTG<br>AATACAAATCACAGAATCGTCGTATGCAGTGAAAACTCTCTTCAATTCTTTATGCCGGTGTTGGGCGCGTTA | 44.76 | 0.22 | 0.71 | -0.06 |

| | | | | | |
|---|---|---|---|---|---|
| | TTTATCGGAGTTGCAGTTGCGCCCGCGAACGACATTTATAATGAACGTGAATTGCTCAACAGTATGAACATT<br>TCGCAGCCTACCGTAGTGTTTGTTTCCAAAAAGGGGTTGCAAAAAATTTTGAACGTGCAAAAAAAATTACCA<br>ATAATCCAGAAAATTATTATCATGGATTCTAAAACGGATTACCAGGGATTTCAGTCGATGTACACGTTCGTC<br>ACATCTCATCTACCTCCCGGTTTTAATGAATACGATTTTGTACCAGAGTCCTTTGATCGTGACAAAACAATT<br>GCACTGATAATGAATTCCTCTGGATCTACTGGGTTACCTAAGGGTGTGGCCCTTCCGCATAGAACTGCCTGC<br>GTCAGATTCTCGCATGCCAGAGATCCTATTTTTGGCAATCAAATCATTCCGGATACTGCGATTTTAAGTGTT<br>GTTCCATTCCATCACGGTTTTGGAATGTTTACTACACTCGGATATTTGATATGTGGATTTCGAGTCGTCTTA<br>ATGTATAGATTTGAAGAAGAGCTGTTTTTACGATCCCTTCAGGATTACAAAATTCAAAGTGCGTTGCTAGTA<br>CCAACCCTATTTTCATTCTTCGCCAAAAGCACTCTGATTGACAAATACGATTTATCTAATTTACACGAAATT<br>GCTTCTGGGGGCGCACCTCTTTCGAAAGAAGTCGGGGAAGCGGTTGCAAAACGCTTCCATCTTCCAGGGATA<br>CGACAAGGATATGGGCTCACTGAGACTACATCAGCTATTCTGATTACACCCGAGGGGGATGATAAACCGGGC<br>GCGGTCGGTAAAGTTGTTCCATTTTTTGAAGCGAAGGTTGTGGATCTGGATACCGGGAAAACGCTGGGCGTT<br>AATCAGAGAGGCGAATTATGTGTCAGAGGACCTATGATTATGTCCGGTTATGTAAACAATCCGGAAGCGACC<br>AACGCCTTGATTGACAAGGATGGATGGCTACATTCTGGAGACATAGCTTACTGGGACGAAGACGAACACTTC<br>TTCATAGTTGACCGCTTGAAGTCTTTAATTAAATACAAAGGATATCAGGTGGCCCCCGCTGAATTGGAATCG<br>ATATTGTTACAACACCCCAACATCTTCGACGCGGGCGTGGCAGGTCTTCCCGACGATGACGCCGGTGAACTT<br>CCCGCCGCCGTTGTTGTTTTGGAGCACGGAAAGACGATGACGGAAAAAGAGATCGTGGATTACGTCGCCAGT<br>CAAGTAACAACCGCGAAAAAGTTGCGCGGAGGAGTTGTGTTTGTGGACGAAGTACCGAAAGGTCTTACCGGA<br>AAACTCGACGCAAGAAAAATCAGAGAGATCCTCATAAAGGCCAAGAAGGGCGGAAAGTCCAAATTGTAA | | | | |
| RL (parental) | ATGGAACAAAAACTCATCTCAGAAGAGGATCTGTCGAGCTCCACTTCGAAAGTTTATGATCCAGAACAAAGG<br>AAACGGATGATAACTGGTCCGCAGTGGTGGGCCAGATGTAAACAAATGAATGTTCTTGATTCATTTATTAAT<br>TATTATGATTCAGAAAAACATGCAGAAAATGCTGTTATTTTTTTACATGGTAACGCGGCCTCTTCTTATTTA<br>TGGCGACATGTTGTGCCACATATTGAGCCAGTAGCGCGGTGTATTATACCAGACCTTATTGGTATGGGCAAA<br>TCAGGCAAATCTGGTAATGGTTCTTATAGGTTACTTGATCATTACAAATATCTTACTGCATGGTTTGAACTT<br>CTTAATTTACCAAAGAAGATCATTTTTGTCGGCCATGATTGGGGTGCTTGTTTGGCATTTCATTATAGCTAT<br>GAGCATCAAGATAAGATCAAAGCAATAGTTCACGCTGAAAGTGTAGTAGATGTGATTGAATCATGGGATGAA<br>TGGCCTGATATTGAAGAAGATATTGCGTTGATCAAATCTGAAGAAGGAGAAAAAATGGTTTTGGAGAATAAC<br>TTCTTCGTGGAAACCATGTTGCCATCAAAAATCATGAGAAAGTTAGAACCAGAAGAATTTGCAGCATATCTT<br>GAACCATTCAAAGAGAAAGGTGAAGTTCGTCGTCCAACATTATCATGGCCTCGTGAAATCCCGTTAGTAAAA<br>GGTGGTAAACCTGACGTTGTACAAATTGTTAGGAATTATAATGCTTATCTACGTGCAAGTGATGATTTACCA<br>AAAATGTTTATTGAATCGGACCCAGGATTCTTTTCCAATGCTATTGTTGAAGGTGCCAAGAAGTTTCCTAAT<br>ACTGAATTTGTCAAAGTAAAAGGTCTTCATTTTTCGCAAGAAGATGCACCTGATGAAATGGGAAAATATATC<br>AAATCGTTCGTTGAGCGAGTTCTCAAAAATGAACAATAA | 36.92 | 0.20 | 0.68 | -0.15 |
| RL-1 | ATGGAACAAAAACTTATTTCAGAGGAGGACCTCTCTTCAAGCACCAGCAAGGTGTACGACCCTGAGCAGCGT<br>AAGAGAATGATCACCGGGCCACAATGGTGGGCCAGATGCAAGCAGATGAATGTGCTGGACAGCTTCATCAAC<br>TATTACGACAGCGAGAAGCACGCCGAGAACGCCGTGATCTTCCTGCACGGCAACGCGGCAAGTAGCTACCTG<br>TGGAGACACGTGGTACCTCACATTGAACCTGTTGCTAGGTGCATAATTCCCGATCTGATTGGAATGGGAAAG | 52.72 | 0.25 | 0.84 | 0.07 |

| | | | | | |
|---|---|---|---|---|---|
| | TCCGGAAAGTCCGGCAATGGAAGCTACAGGCTGCTGGATCATTACAAGTACCTGACCGCTTGGTTCGAGCTG<br>TTGAACCTGCCAAAAAAAATTATTTTTGTTGGACACGATTGGGGCGCCTGCCTTGCCTTCCATTACTCCTAT<br>GAGCACCAAGACAAGATTAAAGCCATTGTGCACGCAGAATCCGTCGTCGACGTGATCGAGAGCTGGGACGAG<br>TGGCCCGACATCGAGGAGGATATTGCCCTGATTAAGAGCGAAGAGGGGGAGAAGATGGTTCTCGAGAACAAC<br>TTCTTCGTGGAAACTATGTTGCCGTCGAAGATTATGAGAAAGCTGGAGCCCGAGGAGTTCGCTGCATACCTC<br>GAGCCCTTCAAGGAGAAGGGTGAGGTCAGGCGGCCAACCCTGTCTTGGCCCCGCGAGATTCCCTTGGTCAAG<br>GGGGGCAAGCCCGACGTGGTCCAGATCGTTCGGAACTACAACGCCTACCTGCGGGCTAGTGACGACCTGCCA<br>AAGATGTTCATTGAGAGCGACCCAGGGTTCTTCTCGAACGCCATCGTGGAGGGGGCGAAGAAATTTCCCAAT<br>ACCGAATTCGTGAAGGTGAAGGGGCTTCACTTCAGCCAGGAGGACGCCCCGGACGAAATGGGCAAGTACATC<br>AAGTCCTTCGTGGAACGCGTACTAAAAAATGAACAATAA | | | | |
| RL-2 | ATGGAGCAGAAACTAATAAGTGAGGAGGATCTCTCTTCTTCTACATCTAAAGTTTACGATCCTGAACAGCGA<br>AAACGAATGATTACAGGGCCTCAGTGGTGGGCGCGCTGTAAACAAATGAATGTTTTGGATTCTTTTATTAAC<br>TACTACGATTCTGAAAAACATGCGGAAAATGCTGTTATCTTTTTGCATGGAAACGCCGCCTCCTCTTACTTA<br>TGGCGTCACGTAGTCCCTCATATTGAACCTGTAGCCCGGTGCATCATTCCTGATTTAATAGGAATGGGCAAA<br>TCTGGAAAATCTGGGAACGGATCTTACAGACTGCTGGATCATTACAAATACTTGACAGCCTGGTTTGAACTG<br>CTAAATTTACCTAAAAAAATTATCTTTGTGGGCCATGATTGGGGGGCCTGCCTTGCCTTTCATTACTCGTAT<br>GAACATCAGGACAAAATCAAAGCAATAGTGCATGCCGAATCAGTGGTTGATGTCATTGAATCCTGGGATGAA<br>TGGCCTGACATTGAAGAAGATATTGCCTTGATCAAAAGTGAGGAAGGAGAAAAAATGGTATTGGAAAACAAC<br>TTCTTTGTAGAGACAATGCTGCCGTCTAAAATTATGCGTAAATTAGAACCTGAAGAATTTGCTGCGTACTTG<br>GAACCTTTTAAAGAAAAAGGGGAAGTTCGTAGACCTACTCTGAGCTGGCCTAGAGAAATTCCACTAGTTAAA<br>GGAGGAAAACCTGATGTCGTGCAGATTGTAAGAAACTACAACGCCTACCTGCGCGCCTCTGATGATTTGCCT<br>AAAATGTTTATTGAATCTGATCCTGGGTTCTTCTCAAATGCCATTGTTGAAGGAGCTAAGAAATTCCCTAAC<br>ACGGAATTTGTGAAAGTGAAAGGACTTCATTTTTCTCAAGAAGATGCTCCAGACGAAATGGGAAAATACATT<br>AAATCCTTTGTCGAGCGCGTATTGAAAAACGAGCAATAA | 40.72 | 0.21 | 0.75 | -0.10 |
| RL-3 | ATGGAGCAGAAGCTGATCAGCGAGGAGGACCTGAGCAGCAGCACCAGCAAGGTGTACGACCCCGAGCAGAGA<br>AAGAGAATGATCACCGGCCCCCAGTGGTGGGCCAGATGCAAGCAGATGAACGTGCTGGACAGCTTCATCAAC<br>TACTACGACAGCGAGAAGCACGCCGAGAACGCCGTGATCTTCCTGCACGGCAACGCCGCCAGCAGCTACCTG<br>TGGAGACACGTGGTGCCCCACATCGAGCCCGTGGCCAGATGCATCATCCCCGACCTGATCGGCATGGGCAAG<br>AGCGGCAAGAGCGGCAACGGCAGCTACAGACTGCTGGACCACTACAAGTACCTGACCGCCTGGTTCGAGCTG<br>CTGAACCTGCCCAAGAAGATCATCTTCGTGGGCCACGACTGGGGCGCCTGCCTGGCCTTCCACTACAGCTAC<br>GAGCACCAGGACAAGATCAAGGCCATCGTGCACGCCGAGAGCGTGGTGGACGTGATCGAGAGCTGGGACGAG<br>TGGCCCGACATCGAGGAGGACATCGCCCTGATCAAGAGCGAGGAGGGCGAGAAGATGGTGCTGGAGAACAAC<br>TTCTTCGTGGAGACCATGCTGCCCAGCAAGATCATGAGAAAGCTGGAGCCCGAGGAGTTCGCCGCCTACCTG<br>GAGCCCTTCAAGGAGAAGGGCGAGGTGAGAAGACCCACCCTGAGCTGGCCCAGAGAGATCCCCCTGGTGAAG<br>GGCGGCAAGCCCGACGTGGTGCAGATCGTGAGAAACTACAACGCCTACCTGAGAGCCAGCGACGACCTGCCC<br>AAGATGTTCATCGAGAGCGACCCCGGCTTCTTCAGCAACGCCATCGTGGAGGGCGCCAAGAAGTTCCCCAAC<br>ACCGAGTTCGTGAAGGTGAAGGGCCTGCACTTCAGCCAGGAGGACGCCCCCGACGAGATGGGCAAGTACATC<br>AAGAGCTTCGTGGAGAGAGTGCTGAAGAACGAGCAGTAA | 60.36 | 0.29 | 1.00 | 0.19 |

| | | | | | |
|---|---|---|---|---|---|
| nanoLuc | ATGGTCTTCACACTCGAAGATTTCGTTGGGGACTGGCGACAGACAGCCGGCTACAACCTGGACCAAGTCCTT<br>GAACAGGGAGGTGTGTCCAGTTTGTTTCAGAATCTCGGGGTGTCCGTAACTCCGATCCAAAGGATTGTCCTG<br>AGCGGTGAAAATGGGCTGAAGATCGACATCCATGTCATCATCCCGTATGAAGGTCTGAGCGGCGACCAAATG<br>GGCCAGATCGAAAAAATTTTTAAGGTGGTGTACCCTGTGGATGATCATCACTTTAAGGTGATCCTGCACTAT<br>GGCACACTGGTAATCGACGGGGTTACGCCGAACATGATCGACTATTTCGGACGGCCGTATGAAGGCATCGCC<br>GTGTTCGACGGCAAAAAGATCACTGTAACAGGGACCCTGTGGAACGGCAACAAAATTATCGACGAGCGCCTG<br>ATCAACCCCGACGGCTCCCTGCTGTTCCGAGTAACCATCAACGGAGTGACCGGCTGGCGGCTGTGCGAACGC<br>ATTCTGGCGTAA | 52.71 | 0.26 | 0.80 | 0.06 |
| dsRED2 | ATGGCCTCCTCCGAGAACGTCATCACCGAGTTCATGCGCTTCAAGGTGCGCATGGAGGGCACCGTGAACGGC<br>CACGAGTTCGAGATCGAGGGCGAGGGCGAGGGCCGCCCCTACGAGGGCCACAACACCGTGAAGCTGAAGGTG<br>ACCAAGGGCGGCCCCCTGCCCTTCGCCTGGGACATCCTGTCCCCCCAGTTCCAGTACGGCTCCAAGGTGTAC<br>GTGAAGCACCCCGCCGACATCCCCGACTACAAGAAGCTGTCCTTCCCCGAGGGCTTCAAGTGGGAGCGCGTG<br>ATGAACTTCGAGGACGGCGGCGTGGCGACCGTGACCCAGGACTCCTCCCTGCAGGACGGCTGCTTCATCTAC<br>AAGGTGAAGTTCATCGGCGTGAACTTCCCCTCCGACGGCCCCGTGATGCAGAAGAAGACCATGGGCTGGGAG<br>GCCTCCACCGAGCGCCTGTACCCCCGCGACGGCGTGCTGAAGGGCGAGACCCACAAGGCCCTGAAGCTGAAG<br>GACGGCGGCCACTACCTGGTGGAGTTCAAGTCCATCTACATGGCCAAGAAGCCCGTGCAGCTGCCCGGCTAC<br>TACTACGTGGACGCCAAGCTGGACATCACCTCCCACAACGAGGACTACACCATCGTGGAGCAGTACGAGCGC<br>ACCGAGGGCCGCCACCACCTGTTCCTGTAG | 63.72 | 0.30 | 0.98 | 0.22 |
| ACTB* | ATGGTGGGCATGGGTCAGAAGGATTCCTATGTGGGCGACGAGGCCCAGAGCAAGAGAGGCATCCTCACCCTG<br>AAGTACCCCATCGAGCACGGCATCGTCACCAACTGGGACGACATGGAGAAAATCTGGCACCACACCTTCTAC<br>AATGAGCTGCGTGTGGCTCCCGAGGAGCACCCCGTGCTGCTGACCGAGGCCCCCCTGAACCCCAAGGCCAAC<br>CGCGAGAAGATGACCCAGATCATGTTTGAGACCTTCAACACCCCAGCCATGTACGTTGCTATCCAGGCTGTG<br>CTATCCCTGTACGCCTCTGGCCGTACCACTGGCATCGTGATGGACTCCGGTGACGGGGTCACCCACACTGTG<br>CCCATCTACGAGGGGTATGCCCTCCCCCATGCCATCCTGCGTCTGGACCTGGCTGGCCGGGACCTGACTGAC<br>TACCTCATGAAGATCCTCACCGAGCGCGGCTACAGCTTCACCACCACGGCCGAGCGGGAAATCGTGCGTGAC<br>ATTAAGGAGAAGCTGTGCTACGTCGCCCTGGACTTCGAGCAAGAGATGGCCACGGCTGCTTCCAGCTCCTCC<br>CTGGAGAAGAGCTACGAGCTGCCTGACGGCCAGGTCATCACCATTGGCAATGAGCGGTTCCGCTGCCCTGAG<br>GCACTCTTCCAGCCTTCCTTCCTGGGCATGGAGTCCTGTGGCATCCACGAAACTACCTTCAACTCCATCATG<br>AAGTGTGACGTGGACATCCGCAAAGACCTGTACGCCAACACAGTGCTGTCTGGCGGCACCACCATGTACCCT<br>GGCATTGCCGACAGGATGCAGAAGGAGATCACTGCCCTGGCACCCAGCACAATGAAGATCAAGATCATTGCT<br>CCTCCTGAGCGCAAGTACTCCGTGTGGATCGGCGGCTCCATCCTGGCCTCGCTGTCCACCTTCCAGCAGATG<br>TGGATCAGCAAGCAGGAGTAG | 55.87 | 0.25 | 0.88 | 0.11 |
| N SARS2 | ATGTCTGATAATGGACCCCAAAATCAGCGAAATGCACCCCGCATTACGTTTGGTGGACCCTCAGATTCAACT<br>GGCAGTAACCAGAATGGAGAACGCAGTGGGGCGCGATCAAAACAACGTCGGCCCCAAGGTTTACCCAATAAT | 47.22 | 0.20 | 0.74 | -0.08 |

| | | | | | |
|---|---|---|---|---|---|
| | ACTGCGTCTTGGTTCACCGCTCTCACTCAACATGGCAAGGAAGACCTTAAATTCCCTCGAGGACAAGGCGTT<br>CCAATTAACACCAATAGCAGTCCAGATGACCAAATTGGCTACTACCGAAGAGCTACCAGACGAATTCGTGGT<br>GGTGACGGTAAAATGAAAGATCTCAGTCCAAGATGGTATTTCTACTACCTAGGAACTGGGCCAGAAGCTGGA<br>CTTCCCTATGGTGCTAACAAAGACGGCATCATATGGGTTGCAACTGAGGGAGCCTTGAATACACCAAAAGAT<br>CACATTGGCACCCGCAATCCTGCTAACAATGCTGCAATCGTGCTACAACTTCCTCAAGGAACAACATTGCCA<br>AAAGGCTTCTACGCAGAAGGGAGCAGAGGCGGCAGTCAAGCCTCTTCTCGTTCCTCATCACGTAGTCGCAAC<br>AGTTCAAGAAATTCAACTCCAGGCAGCAGTAGGGGAACTTCTCCTGCTAGAATGGCTGGCAATGGCGGTGAT<br>GCTGCTCTTGCTTTGCTGCTGCTTGACAGATTGAACCAGCTTGAGAGCAAAATGTCTGGTAAAGGCCAACAA<br>CAACAAGGCCAAACTGTCACTAAGAAATCTGCTGCTGAGGCTTCTAAGAAGCCTCGGCAAAAACGTACTGCC<br>ACTAAAGCATACAATGTAACACAAGCTTTCGGCAGACGTGGTCCAGAACAAACCCAAGGAAATTTTGGGGAC<br>CAGGAACTAATCAGACAAGGAACTGATTACAAACATTGGCCGCAAATTGCACAATTTGCCCCCAGCGCTTCA<br>GCGTTCTTCGGAATGTCGCGCATTGGCATGGAAGTCACACCTTCGGGAACGTGGTTGACCTACACAGGTGCC<br>ATCAAATTGGATGACAAAGATCCAAATTTCAAAGATCAAGTCATTTTGCTGAATAAGCATATTGACGCATAC<br>AAAACATTCCCACCAACAGAGCCTAAAAAGGACAAAAAGAAGAAGGCTGATGAAACTCAAGCCTTACCGCAG<br>AGACAGAAGAAACAGCAAACTGTGACTCTTCTTCCTGCTGCAGATTTGGATGATTTCTCCAAACAATTGCAA<br>CAATCCATGAGCAGTGCTGACTCAACTCAGGCCTAAT | | | | |
| N1 SARS2 | ATGAGCGACAATGGCCCCCAGAACCAGAGAAACGCTCCTCGCATCACCTTCGGCGGCCCATCTGATAGCACC<br>GGGAGCAATCAGAATGGCGAGAGGAGCGGGGCCAGATCAAAACAGAGGAGACCCCAGGGGCTGCCAAACAAT<br>ACCGCCAGCTGGTTTACTGCCCTGACTCAGCACGGCAAGGAGGATCTGAAGTTCCCTAGGGGTCAGGGCGTG<br>CCTATCAATACAAACTCTAGCCCCGATGACCAGATCGGATACTACCGCCGCGCTACACGGAGAATCAGGGGC<br>GGCGATGGAAAAATGAAAGACCTGTCTCCCCGGTGGTACTTCTACTATCTGGGCACCGGCCCTGAAGCTGGA<br>CTTCCCTATGGTGCCAACAAGGACGGAATCATTTGGGTGGCCACCGAAGGCGCCCTGAATACACCAAAGGAC<br>CACATCGGCACCAGGAATCCTGCTAACAATGCTGCAATCGTGCTGCAGCTGCCCCAGGGAACTACCCTGCCT<br>AAGGGTTTCTACGCTGAAGGCTCCCGCGGGGGCTCCCAGGCCTCCAGCAGGTCTTCCAGCAGATCCCGCAAT<br>TCCTCCCGCAATAGCACCCCCGGCTCCTCTCGGGGCACCAGCCCAGCCAGGATGGCTGGAAATGGCGGCGAC<br>GCCGCTCTTGCCCTGCTGCTGCTGGACAGGCTGAATCAGCTGGAGTCTAAGATGAGCGGGAAGGGCCAGCAG<br>CAGCAGGGCCAGACCGTGACCAAGAAGTCCGCAGCCGAGGCCAGCAAGAAGCCCAGGCAGAAAAGAACAGCC<br>ACAAAAGCCTACAACGTCACTCAGGCCTTTGGCAGGAGGGGACCCGAACAGACTCAGGGCAACTTTGGCGAC<br>CAGGAGCTGATCCGCCAGGGAACCGACTACAAGCACTGGCCTCAGATCGCCCAGTTCGCCCCCTCTGCCAGC<br>GCTTTTTTTGGCATGAGCAGGATCGGAATGGAGGTGACTCCAAGCGGCACCTGGCTGACTTACACCGGGGCT<br>ATTAAGCTGGACGACAAAGATCCCAACTTCAAGGATCAGGTGATCCTCCTGAACAAGCACATCGACGCCTAC<br>AAGACCTTCCCCCCTACCGAGCCTAAGAAGGATAAGAAGAAAAAGGCCGACGAGACCCAGGCCCTCCCTCAG<br>AGACAGAAAAAGCAGCAGACCGTGACCCTGCTGCCTGCCGCCGATCTGGACGATTTCTCTAAACAGCTGCAG<br>CAGAGCATGAGTTCCGCCGACAGTACCCAGGCCTGA | 59.00 | 0.25 | 0.92 | 0.08 |
