## Supplementary table 2 for "The differential effect of SARS-COV-2 NSP1 on mRNA translation and stability reveals new insights linking ribosome recruitment, codon usage and virus evolution"

| 5' UTR | sequence | reporter |
|---|---|---|
| 94 | AGATCCGCTAGCGCTACCGGACTCAGATCTCGAGCTCAAGCTTCGAATTCTGCAGTCGACGGTACCGCGGGCCC<br>GGGATCCACCGGTCGCCACC**ATG** | EGFP,dsRed2,<br>nanoLuc,N<br>SARS-CoV-2 |
| 82 | AGATCCGCTAGCGCTACCGGACTCAGATCTCGAGCTCAAGCTTCGAATTCTGCAGTCGACGGTACCGCGGGCCC<br>GGGATCCA**ATG** | FFL, RL |
| 223 | AGCGCTACCGGACTCAGAATCTCGAGCTCAAGCTTCGAATTCTGCAGTCGACGGTACCGCGGGCCCCTTGGCCT<br>GGCTTCGCGCTACGCCGGCGCGCGCGGCCGCGAATTGTAGGTGGGCGTGGCCTCCAAGGGCGTGGCGGCATT<br>CGTGGTCTCCATCGCCTGCCATAAAACACTTGTGTTGGTAGGAAATCCATAGAGCGCCCCTATAGTGGGGATCC<br>A**ATG** | RL |
| 424 | AGATCCGCTAGCGCTACCGGACTCAGATCTCGAGCTCTCTGGCTAACTAGGGAACCCACTGCTTAAGCCTCAAT<br>AAAGCTTGCCTTGAGTGCTTCAAGTAGTGTGTGCCCGTCTGTTGTGTGACTCTGGTAACTAGAGATCCCTCAGA<br>CCCTTTTAGTCAGTGTGGAAAATCTCTAGCAGTGGCGCCCGAACAGGGACTTGAAAGCGAAAGGGAAACCAGAG<br>GAGCTCAAGCTTCGAATTCTGCAGTCGACGGTACCGCGGGCCCCTTGGCCTGGCTTCGCTCTACGCCGGCGCGC<br>GCGCGGCGCGAATTGTAGGTGGGCGTGGCCTCCAAGGGCGTGGCGGCATTCGTGGTCTCCATCGCCTGCCATAA<br>AACACTTGTGTTGGTAGGAAATCCATTCTAGAGCGCCCCTATAGTGGGGATCCA**ATG** | RL |
| 626 | AGATCTCGAGCTCAAGCTTCGAATTCTGCAGTCGACGGTACCGCGGGCCCAGGTAGACAATATTACACCTGTCC<br>TACTGGCATTGAGAACTTTTGCCCAGAGCAAAAGAGCATTCCAAGCCATCAGAGGGGAAATAAAGCATCTCTAC<br>GGTGGTCCTAAATAGTCAGCATAGTACATTTCATCTGACTAATACTACAACACCACCACCTCTAGCGCTACCGG<br>ACTCAGATCTCGAGCTCTCTGGCTAACTAGGGAACCCACTGCTTAAGCCTCAATAAAGCTTGCCTTGAGTGCTT<br>CAAGTAGTGTGTGCCCGTCTGTTGTGTGACTCTGGTAACTAGAGATCCCTCAGACCCTTTTAGTCAGTGTGGAA<br>AATCTCTAGCAGTGGCGCCCGAACAGGGACTTGAAAGCGAAAGGGAAACCAGAGGAGCTCAAGCTTCGAATTCT<br>GCAGTCGACGGTACCGCGGGCCCCTTGGCCTGGCTTCGCTCTACGCCGGCGCGCGCGGCGCGAATTGTAGGT<br>GGGCGTGGCCTCCAAGGGCGTGGCGGCATTCGTGGTCTCCATCGCCTGCCATAAAACACTTGTGTTGGTAGGAA<br>ATCCATTCTAGAGCGCCCCTATAGTGGGGATCCA**ATG** | RL |
| SL20 | AGATCCGCTAGCGCTACCGGACTCAGATCTCGAGCTCAAGCTTCGAATTCTGCAGTCGACGGTACCGCGGGCCC<br>CGACCCGGGCCCGCGGTACGCCGATAGGCGTACGGGATCCA**ATG** | FFL,RL |
| SL50 | AGATCCGCTAGCGCTACCGGACTCAGATCTCGAGCTCAAGCTTCGAATTCTGCAGTCGACCCGGGCCCGCGGAG<br>TACTCCGCGGGCCCGCGGAGTACCGCGGGCCCGGGATCCA**ATG** | FFL,RL |
| TISU | AGATCAAG**ATG** | EGFP,RL |
| Gless | AGCCACTATCTCACACCTTTCCTCACTCTTTCCTCACACTTCTTTCTACACTCTTCACAAAAAATAATTTCTCA | RL,RL3 |

| | | |
|---|---|---|
| | CTTCCTATTCTTCTCCCCCATCCCTCATTCCTCAATCATTCCTTCCCCATTCACTTCAATCATTCCAA**ATG** | |
| Gless-SL1 | AGAAAAGATAGTGGAAACCACTATCTCACACCTTTCCTCACTCTTTCCTCACACTTCTTTCTACACTCTTCACA AAAAATAATTTCTCACTTCCTATTCTTCTCCCCCATCCCTCATTCCTCAATCATTCCTTCCCATCCAA**ATG** | RL |
| Gless-SL3 | AGCCACTATCTCACACCTTTCCTCACTCTTTCCTCACACTTCTTTCTACACTCTTCACAAAAAATAATTTCTCA CTTCCTATTCTTCTCCCCCATCCCTCATTCCTCAATCATTCCGAGATAGTGGAAACCACTATCTCCAA**ATG** | RL |
| Gless-M1 | AGCCACGAGCTCGGAGCTTGCCGCACTCGGTCCTCACACTTCTTTCTACACTCTTCACAAAAAATAATTTCTCA CTTCCTATTCTTCTCCCCCATCCCTCATTCCTCAATCATTCCTTCCCCATTCACTTCAATCATTCCAA**ATG** | RL |
| Gless-M2 | AGCCACTATCTCACACCTTTCCTCACTCTTTCCTCACACTTCTTTCTACACTCTTCACAAAAAATAATTTCTCA CTTCCTATTCTTCTCCCCCATCCCTCATTCCTCAATCATGGCGACGCCATTGACTTTGATCATTCCAA**ATG** | RL |
| Gless-M3 | AGCCACGAGCTCGGAGCTTGCCGCACTCGGTCCTCACACTTCTTTCTACACTCTTCACAAAAAATAATTTCTCA CTTCCTATTCTTCTCCCCCATCCCTCATTCCTCAATCATGGCGACGCCATTGACTTTGATCATTCCAA**ATG** | RL |
| Gless-M4 | AGCCACGAGAGAGAGAGATTCCTCACTCTTTCCTCACACTTCTTTCTACACTCTTCACAAAAAATAATTTCTCA CTTCCTATTCTTCTCCCCCATCCCTCATTCCTCAATCATTCCTTCCCCATTCACTTCAATCATTCCAA**ATG** | RL |
| Slot | AGCACAACAACAACAAACCCTCGAACAACAACAACAACAACAACAACACC**ATG** | RL |
| Gless_8 | AGCCACAA**ATG** | RL |
| Gless_12 | AGCCACTTCCAA**ATG** | RL |
| Gless_15 | AGCCACTATTTCCAA**ATG** | RL |
| Gless_22 | AGCCACTATCTCATCATTCCAA**ATG** | RL |
| Gless_30 | AGCCACTATCTCACACCTCAATCATTCCAA**ATG** | RL |

| | | |
|---|---|---|
| Gless_67 | AGCCACTATCTCACACCTTTCCTCACTCTTTCCTCCATTCCTTCCCCATTCACTTCAATCATTCCAA**ATG** | RL |
| 5'L-N | ATTAAAGGTTTATACCTTCCCAGGTAACAAACCAACCAACTTTCGATCTCTTGTAGATCTGTTCTCTAAACGAACAAACT**ATG** | RL, N SARS2 |
| 5'L(Gs)-N | ATTAAAGGTGGAGACGAGCCGAGGTAACAAACCAACCAACTTTCGATCTCTTGTAGATCTGTTCTCTAAACGAACAAACT**ATG** | RL, N SARS2 |
| Gless-uATG_8 | AGCCACTATGCACACCTTTCCTCACTCTTTCCTCACACTTCTTTCTACACTCTTCACAAAAAATAATTTCTCACTTCCTATTCTTCTCCCCCATCCCTCATTCCTCAATCATTCCTTCCCCATTCACTTCAATCATTCCAA**ATG** | RL |
| Gless-uATG_13 | AGCCACTATATCATGCCCTTTCCTCACTCTTTCCTCACACTTCTTTCTACACTCTTCACAAAAAATAATTTCTCACTTCCTATTCTTCTCCCCCATCCCTCATTCCTCAATCATTCCTTCCCCATTCACTTCAATCATTCCAA**ATG** | RL |
| Gless-uATG_18 | AGCCACTATCTCACACCATGCCTCACTCTTTCCTCACACTTCTTTCTACACTCTTCACAAAAAATAATTTCTCACTTCCTATTCTTCTCCCCCATCCCTCATTCCTCAATCATTCCTTCCCCATTCACTTCAATCATTCCAA**ATG** | RL |
| Gless-uATG_28 | AGCCACTATCTCACACCTTTCCTCACTATGATCCTCACACTTCTTTCTACACTCTTCACAAAAAATAATTTCTCACTTCCTATTCTTCTCCCCCATCCCTCATTCCTCAATCATTCCTTCCCCATTCACTTCAATCATTCCAA**ATG** | RL |
| Gless-uATG_112 | AGCCACTATCTCACACCTTTCCTCACTCTTTCCTCACACTTCTTTCTACACTCTTCACAAAAAATAATTTCTCACTTCCTATTCTTCTCCCCCATCCCTCATTCCTCAATCATGCCTTCCCCATTCACTTCAATCATTCCAA**ATG** | RL |
| Gless-M4-uATG_8 | AGCCACGATGAAGAGAGAGATTCCTCACTCTTTCCTCACACTTCTTTCTACACTCTTCACAAAAAATAATTTCTCACTTCCTATTCTTCTCCCCCATCCCTCATTCCTCAATCATTCCTTCCCCATTCACTTCAATCATTCCAA**ATG** | RL |
| Gless-M4-uATG_28 | AGCCACGAGAGAGAGAGATTCCTCACTATGATCCTCACACTTCTTTCTACACTCTTCACAAAAAATAATTTCTCACTTCCTATTCTTCTCCCCCATCCCTCATTCCTCAATCATTCCTTCCCCATTCACTTCAATCATTCCAA**ATG** | RL |
| G-low | AGATCCGGCTTATTTCCTCCTTCTCCACTATCTCACACCTTTCCTCACTCTTTCCTCACACTTCTTTCTACACTCTTCACAAAAAATAATTTCTCACTTCCTATTCTTCTCCCCCATCCCTCATTCCTCAATCATTCCTTCCCCATTCACTTCAATCATTCCAAGGATCCA**ATG** | FFL, RL, RL3, EGFP |
