## Supplementary table 3 for "The differential effect of SARS-COV-2 NSP1 on mRNA translation and stability reveals new insights linking ribosome recruitment, codon usage and virus evolution"

| oligo name | oligo sequence | purpose |
|---|---|---|
| 5'-EX-NSP1 | CACAATCTCGAGAATTCATGGAGAGCCTGGTGCCTGGCTTCAACG | cloning |
| 3'-NB-NSP1 | CACACGGATCCGCGGCCGCTTAGCCACCGTTCAGTTCACGCATCAGC | cloning |
| 5'-N-hACTB | GACGACCCATGGTGGGCATGGGTCAGAAGGATTCC | cloning |
| 3'-X-hACTB | CACCACATCTAGACTAGAAGCATTTGCGGTGGACGATGGA | cloning |
| 5'-161 pEGFPN1 | GGGACTTTCCATTGACGTCAATGGGTGG | cloning |
| 3'-TISU-SacI | CTGGTTGCGGAGCTCACCGCCGCCATCTTGATCTGACGGTT | cloning |
| 5'-NanoL | CGACAGGATCCACCATGGTCTTCACACTCGAAGATTTC | cloning |
| 3'-NanoL | GTGTGTGCGGCCGCTACGCCAGAATGCGTTCGCAC | cloning |
| 5'-18S_9 | CGGGGAGGTAGTGACG | RT-PCR 18S |
| 3'-18S_4 | CGGTATCCAGGCGGCTCGG | RT-PCR 18S |
| 5'-EGFP | GAACCGCATCGAGCTGAA | RT-PCR EGFP |
| 3'-EGFP | TGCTTGTCGGCCATGATATAG | RT-PCR EGFP |
| 5'-RL | GTTAGGAATTATAATGCTTATCTACG | RT-PCR RL |
| 3'-RL | CTTGGCACCTTCAACAATAGC | RT-PCR RL |
| 5'-RL3 | GGAGAAGGGCGAGGTGAGAAG | RT-PCR RL3 |
| 3'-RL3 | CGATGGCGTTGCTGAAGAAG | RT-PCR RL3 |
| 5'-NanoL_qPCR | GGAGGTGTGTCCAGTTTGTT | RT-PCR nanoL |
| 3'-NanoL_qPCR | ATGTCGATCTTCAGCCCATTT | RT-PCR nanoL |
| 5'-dsRed | GAACGTCATCACCGAGTTCA | RT-PCR dsRed |
| 3'-dsRed | GGGTGCTTCACGTACACCTT | RT-PCR dsRed |
| 5'-N-SARS2 | AAGCTGGACTTCCCTATGGTG | RT-PCR N SARS2 |
| 3'-N-SARS2 | CGATTGCAGCATTGTTAGCAGG | RT-PCR N SARS2 |
